# SLC25A39 is necessary for mitochondrial glutathione import in mammalian cells

**DOI:** 10.1101/2021.09.15.460381

**Authors:** Ying Wang, Frederick S. Yen, Xiphias Ge Zhu, Rebecca C. Timson, Ross Weber, Changrui Xing, Yuyang Liu, Benjamin Allwein, Hanzhi Luo, Hsi-Wen Yeh, Søren Heissel, Gokhan Unlu, Eric R. Gamazon, Michael G. Kharas, Richard Hite, Kivanç Birsoy

## Abstract

Glutathione (GSH) is a small molecule thiol abundantly present in all eukaryotes with key roles in oxidative metabolism^1^. Mitochondria, as the major site of oxidative reactions, must maintain sufficient levels of GSH to perform protective and biosynthetic functions^2^. GSH is exclusively synthesized in the cytosol, yet the molecular machinery involved in mitochondrial GSH import remain elusive. Here, using organellar proteomics and metabolomics approaches, we identify SLC25A39, a mitochondrial membrane carrier of unknown function, to regulate GSH transport into mitochondria. SLC25A39 loss reduces mitochondrial GSH import and abundance without impacting whole cell GSH levels. Cells lacking both SLC25A39 and its paralog SLC25A40 exhibit defects in the activity and stability of iron-sulfur cluster containing proteins. Moreover, mitochondrial GSH import is necessary for cell proliferation *in vitro* and red blood cell development in mice. Remarkably, the heterologous expression of an engineered bifunctional bacterial GSH biosynthetic enzyme (*GshF*) in mitochondria enabled mitochondrial GSH production and ameliorated the metabolic and proliferative defects caused by its depletion. Finally, GSH availability negatively regulates SLC25A39 protein abundance, coupling redox homeostasis to mitochondrial GSH import in mammalian cells. Our work identifies SLC25A39 as an essential and regulated component of the mitochondrial GSH import machinery.

## Main

Glutathione, a tripeptide with an active thiol, is the predominant antioxidant in eukaryotic cells, existing in millimolar concentrations^4^. While GSH is commonly recognized as a component of antioxidant defense systems, it has wide range of roles in disulfide bond formation, detoxification of xenobiotics, metabolite transport, formation of iron-sulfur clusters and cellular signaling. Most eukaryotic cells synthesize glutathione by the consecutive actions of γ-glutamate-cysteine ligase (GCL) and glutathione synthetase (GS), enzymes localized in the cytosol. Despite its exclusive cytosolic production^5^, GSH is also abundantly present in many organelles including peroxisomes, the nucleus, ER and mitochondria^6–8^. In particular, as the most redox active organelle, mitochondria contain 10-15% of total cellular GSH^9^. Because mitochondria lack GSH biosynthetic machinery^5^ and GSH is negatively charged under physiological conditions^10^, it must enter mitochondria through a dedicated transport process. Indeed, previous studies using isolated mitochondria supported for the existence of a mitochondrial glutathione transporter^11,12^, but the molecular machinery involved in mitochondrial GSH transport has remained elusive.

### Global analysis of the mitochondrial proteome in response to cellular GSH depletion

To identify mitochondrial proteins regulated by GSH availability, we used quantitative proteomics and analyzed mitochondria immunopurified from HeLa cells expressing a mitotag (3XHA-OMP25-mCherry) grown in standard media or treated with the glutathione synthesis inhibitor, buthionine sulfoximine (BSO)^13^. BSO treatment decreases GSH levels in whole cells as well as in purified mitochondria with similar kinetics^14^ (Fig.1a). Consistent with the robust mitochondrial immunopurification^15^, peptides identified from mitochondrial fractions were significantly enriched for proteins annotated as mitochondrial (Extended Data Fig. 1a). After BSO treatment compared to untreated cells, 296 proteins exhibited significant alterations in abundance (p-value<0.01, change in abundance>1.5) (Fig 1b,c; Extended Data Table 1). Gene ontology analysis revealed that many components of the mitochondrial translation machinery are downregulated in response to GSH depletion, a phenotype previously linked to electron transport chain dysfunction^14^ (Fig. 1c). GSH depletion also activates the nuclear factor erythroid 2-related factor 2 (*NRF2*), a transcription factor that induces expression of several cellular antioxidant defense genes^16,17^. Indeed, two of the top upregulated proteins, ALAS1 and HMOX1 are downstream mitochondrial targets of the transcription factor NRF2 (Fig. 1c). In individual immunoblotting experiments, we confirmed enrichment of mitochondrial markers and multiple proteins upregulated in response to BSO as early as 1 day after treatment (Fig. 1d). Among the most upregulated proteins upon GSH depletion was SLC25A39, an uncharacterized mitochondrial membrane transporter (Fig. 1d, Extended Data Fig. 1b). Induction of SLC25A39 protein levels was generalizable as two additional cell lines from distinct tissue origins (K562 and HEK-293T) displayed similar responses to BSO (Fig. 1e). Furthermore, an antibody targeting endogenous SLC25A39 immunohistochemically colocalized with the inner mitochondrial membrane protein ATP5A in HeLa cells in response to BSO treatment (Fig. 1f). Similar to pharmacologic inhibition of GSH synthesis, loss of GCLC, the rate limiting enzyme of GSH synthesis, also strongly induced SLC25A39 protein (Fig. 1g). Given that SLC25A39 was the only mitochondrial membrane transporter regulated by GSH depletion and its unknown function, we focused our attention on SLC25A39.

**Fig. 1:**
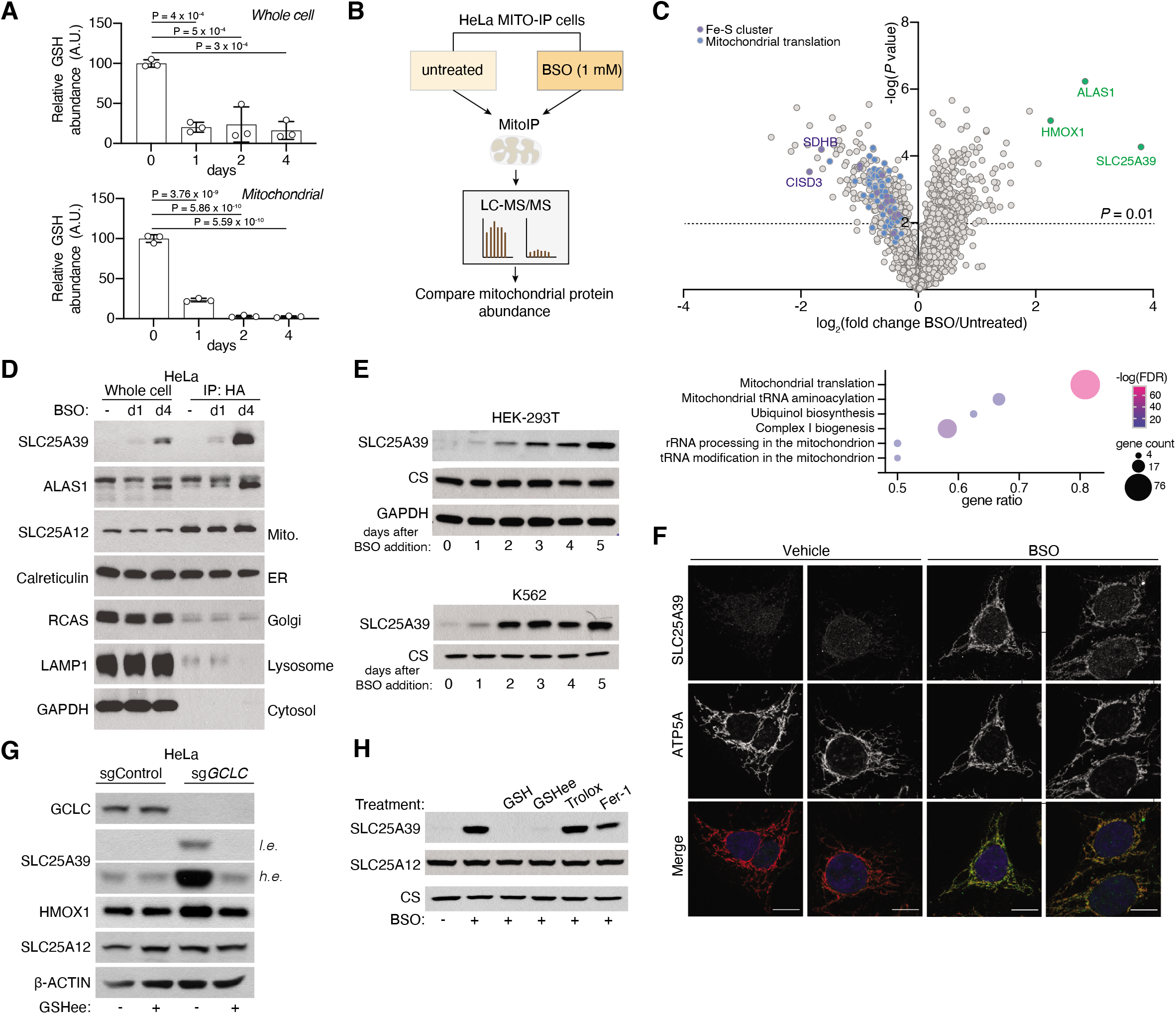
Global analysis of mitochondrial proteome in response to cellular GSH depletion. A. Relative abundance of GSH in whole cells (top) and mitochondria (bottom) immune isolated from HeLa cells treated with BSO (1 mM) for the indicated times compared to untreated controls. Bars represent mean ± s.d.; *n* = 3 biologically independent samples. Statistical significance was determined by one-way ANOVA followed by Bonferroni post-hoc analysis. B. Schematics of proteomics analysis of mitochondria isolated from indicated HeLa cells. C. Volcano plot showing relative fold change (log_2_) in mitochondrial protein abundance vs P values (-log) from HeLa cells treated with BSO (1 mM) for 4 days compared to untreated (top). The dotted line represents *P*=0.01. Colors denote indicated protein families. Gene ontology analysis of proteins altered by BSO treatment (bottom). D. Immunoblot analysis of indicated proteins in whole-cell lysates and mitochondria isolated from HeLa cells treated with BSO (1 mM) for the indicated days. E. Immunoblot of indicated proteins in the HEK-293T (top) and K-562 (bottom) cells treated with BSO (1 mM;10 μM, respectively) for the indicated days. CS and GAPDH were used as loading controls. F. Immunofluorescence analysis of indicated proteins in HeLa cells treated with vehicle or BSO (1 mM) after 24 hours. Micrographs are representative images. Scale bar, 10 μm. G. Immunoblot of indicated proteins in the HeLa cells infected with the indicated sgRNAs. Cells were plated 6 days after transduction and treated for 4 days with GSHee (10 mM). SLC25A12 and β-ACTIN were used as loading controls. H. Immunoblot of indicated proteins in the HeLa cells treated with BSO (1 mM) and co-treated with either GSH (10 mM), GSHee (10 mM), Trolox (50 μM) or Ferrostatin-1 (5 μM) for 48 hours. SLC25A12 and CS were used as loading controls.

We first sought to determine the basis for SLC25A39 protein regulation by BSO. Interestingly, BSO treatment did not affect SLC25A39 mRNA levels (Extended Data Fig 1c), but could induce SLC25A39 protein accumulation when an SLC25A39 cDNA lacking UTR elements was expressed ectopically under a viral promoter (Extended Data Fig 1d), thereby suggesting regulation at the post-translational level. We next asked whether reactive oxygen species (ROS) or subsequent NRF2 induction caused by GSH depletion could impact SLC25A39 levels. Neither increasing ROS or lipid peroxides by treating cells with hydrogen peroxide or ferroptosis inducers (RSL-3), nor activating NRF2 by small molecules (KI696) impacts SLC25A39 protein levels (Extended Data Fig 1e-g). These results indicate that SLC25A39 protein levels are regulated independently of NRF2 activation or oxidative stress. Consistent with these findings, supplementation of only GSH/GSHee, but not other antioxidants (Trolox, Ferrostatin-1) prevented induction in SLC25A39 protein levels upon GSH depletion (Fig. 1g,h, Extended Data Fig. 1h). Collectively, these results suggest that SLC25A39 protein abundance is directly regulated by cellular glutathione availability.

### SLC25A39 mediates mitochondrial GSH import

As SLC25A39 is predicted to be a member of the SLC25A family of mitochondrial small molecule transporters^18^ and is regulated by GSH availability, we hypothesized that it may be involved in glutathione metabolism through its presumed transport activity. To address this in an unbiased way, we profiled mitochondrial metabolites of SLC25A39 knockout cell lines and those complemented with SLC25A39 cDNA by LC-MS. Of note, we performed this analysis in 3 independent human cell lines expressing mitotags that enable immunopurification of mitochondria and identification of mitochondrial metabolites whose abundance changes upon SLC25A39 loss^15^. Remarkably, among all detected metabolites, we observed the largest reductions in the levels of oxidized or reduced glutathione in mitochondria of SLC25A39 knockout cells (Fig. 2a, Extended Data Fig. 2a,b). Loss of SLC25A39 did not impact the levels of whole cell glutathione or most other mitochondrial metabolites but caused a specific 5-10 fold reduction in that of mitochondrial glutathione (Fig. 2b, Extended Data Fig 2c). These results suggest that SLC25A39 is required specifically for the maintenance of mitochondrial GSH pools.

**Fig. 2:**
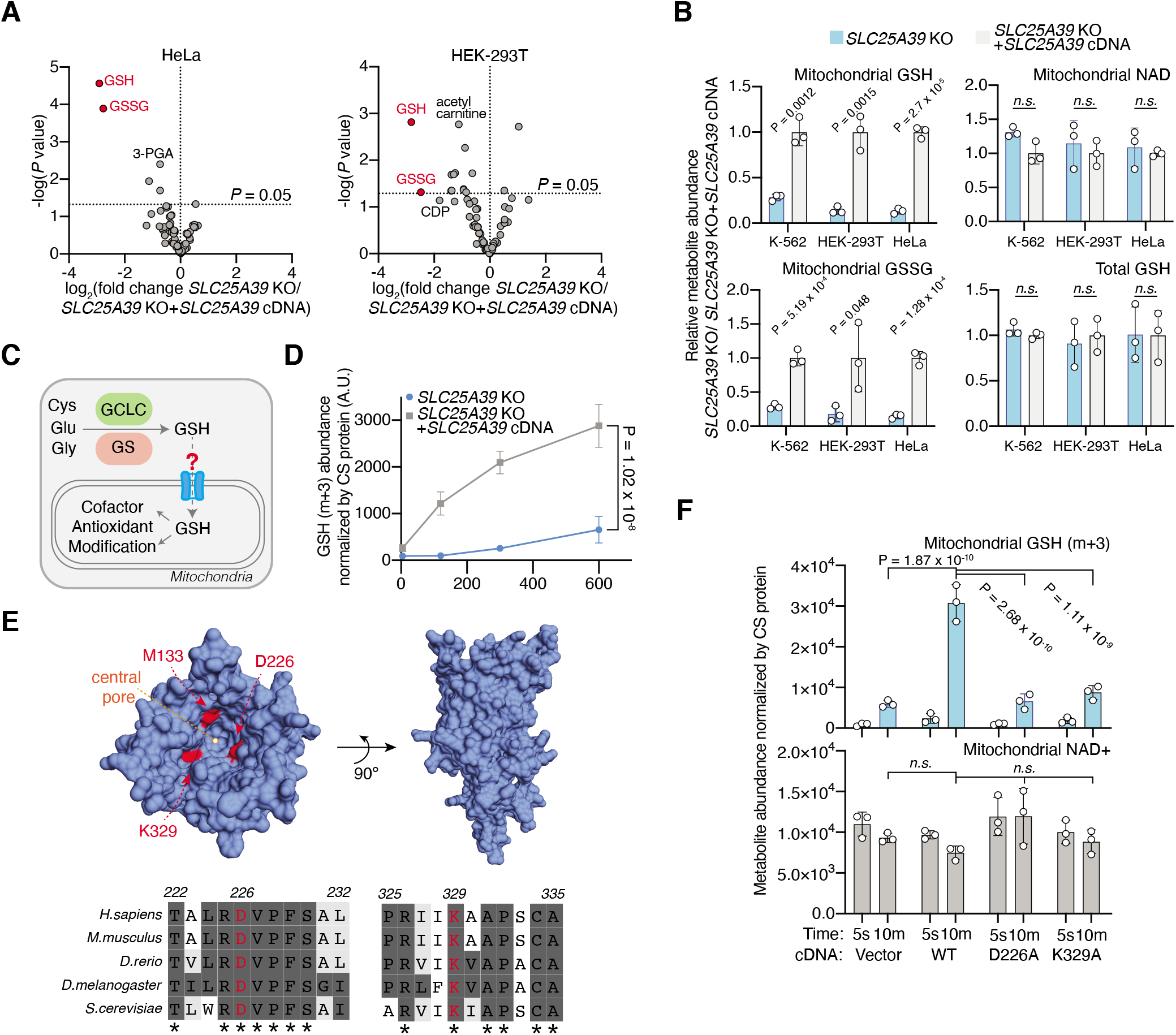
SLC25A39 mediates mitochondrial GSH import. A. Volcano plots showing the fold change in mitochondrial metabolite abundance (log_2_) vs *P* values (-log) from indicated SLC25A39 knockout cell lines expressing a vector control or *SLC25A39* cDNA. Red data points highlight GSH and GSSG. The dotted line represents *P*=0.05. B. Relative metabolite abundance of indicated whole cell or mitochondrial metabolites from K-562, HEK-293T and HeLa SLC25A39 knockout cells lines compared to those expressing *SLC25A39* cDNA. C. Schematic of GSH synthesis in the cytosol and its transport into mitochondria. Cys, cysteine; Gly, glycine; Glu, glutamine; GCLC, Glutamate-cysteine ligase, GS: glutathione synthetase. D. Uptake of [^13^C_2_,^15^N]-GSH into mitochondria isolated from *SLC25A39* knockout HEK-293T cells expressing a vector control or *SLC25A39* cDNA during the indicated times. Metabolite abundance is normalized by CS protein levels. E. Structural and sequence comparison of SLC25A39 with the ADP/ATP carrier identifies methionine 133 (M133), aspartic acid 226 (D226) and lysine 329 (K329) as potential substrate contact points (top). Sequence alignment of SLC25A39 homologs for indicated residues (bottom). F. Uptake of [^13^C_2_,^15^N]-GSH into mitochondria isolated from SLC25A39 knockout HEK-293T cells expressing a vector control or indicated SLC25A39 cDNAs at 5 seconds and 10 minutes. Metabolite abundance is normalized by CS protein levels. Mitochondrial NAD+ abundance is shown to indicate the mitochondrial health (bottom). **b**, **d**, **f**, Bars represent mean ± s.d.; **a, b, d, f,** *n* = 3 biologically independent samples. Statistical significance was determined by two-tailed unpaired *t*-test for **b and** two-way ANOVA followed by Bonferroni post-hoc analysis for **d** and **f**.

Given the strong decrease in the steady state levels of mitochondrial GSH upon SLC25A39 loss, we reasoned that SLC25A39 may mediate GSH import into mitochondria. Previous studies have shown isolated mitochondria are able to import glutathione^11^, but the transporter machinery responsible for this activity has remained elusive. To test this possibility, we established and performed isotope labeled glutathione (GSH-(glycine-^13^C_2_,^15^N)) uptake assays using mitochondria isolated from SLC25A39 knockout cells and those expressing an *SLC25A39* cDNA (Extended Data Fig. 2d,e). During a 10 min uptake assay, mitochondria containing SLC25A39 took up 6-fold more glutathione than those lacking SLC25A39 (Fig. 2d). To determine whether SLC25A39 transport function is essential for GSH import into mitochondria, we generated a homology model of SLC25A39 based on that of the bovine ADP/ATP transporter^19^ that facilitated the identification of three potential substrate binding residues, K329, M133 and D226 (Fig. 2e). Due to their high conservation among SLC25A39 homologs across multiple species, we focused on D226 and K329; and expressed these mutant cDNAs in SLC25A39 knockout HEK-293T cells (Fig. 2f). Remarkably, mutation of each residue to alanine completely abolished the uptake of glutathione, without impacting the levels of NAD+ (Fig. 2f). These data suggest that SLC25A39 mediates GSH import into mitochondria of mammalian cells.

### Mitochondrial GSH import by SLC25A39/40 is essential for cell proliferation

Because SLC25A39 loss did not impact cell proliferation and did not completely abolish mitochondrial GSH import (Fig. 2d), we hypothesized that other metabolic genes or transporters may overcome the decrease in mitochondrial GSH levels. To test this possibility, we performed CRISPR-Cas9 based genetic screens in both Jurkat and HEK-293T cells using a metabolism focused sgRNA library^20^ and searched for genes essential for cellular proliferation in the absence of SLC25A39 (Fig. 3a, Extended Data Table 2). Remarkably, a top scoring gene from both screens was SLC25A40, the mitochondrial SLC25A family transporter with the highest sequence homology to SLC25A39 (Fig 3b,c; Extended Data Fig. 3a, b). Interestingly, a co-evolution analysis for SLC25A39 across a large set of eukaryotes also retrieved SLC25A40 as the top scoring gene (z score= 9.86), indicating a functional association between these transporters (Extended Data Fig. 3c, d). Consistent with the screening results, cells lacking both SLC25A39 and SLC25A40 were unable to proliferate under standard culture conditions (Fig 3d, Extended Data Fig. 3e). Furthermore, overexpressing SLC25A40 in SLC25A39 knockout cells was sufficient to restore mitochondrial GSH levels, suggesting its redundant function in enabling mitochondrial GSH import alongside SLC25A39 (Fig. 3e, Extended Data Fig. 3f). Notably, also scoring in HEK-293T cells was Thioredoxin Reductase 2 (TXNRD2), a component of the thioredoxin (Trx) dependent antioxidant system specifically localized to mitochondria (Fig 3c; Extended Data Fig. 3a). This is in line with the presence of the two parallel thiol antioxidant system based on GSH or thioredoxin-2 in mitochondria^21,22^.

**Fig. 3:**
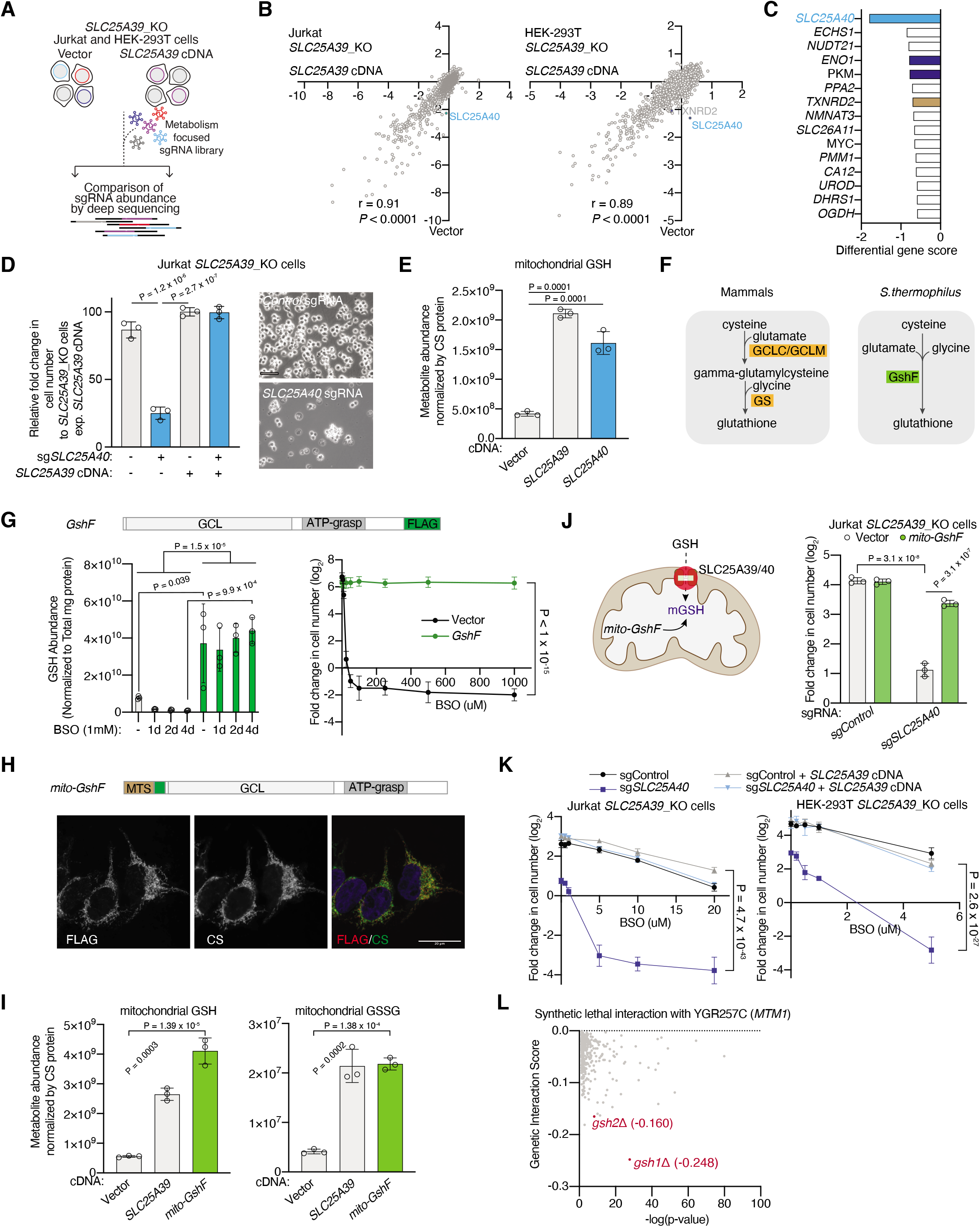
Mitochondrial GSH import by SLC25A39/40 is essential for cell proliferation. A. Schematic depicting the metabolism focused CRISPR genetic screens in Jurkat and HEK-293T SLC25A39 knockout cells expressing a vector control or *SLC25A39* cDNA. B. CRISPR genes scores in indicated Jurkat (left) and HEK-293T (right) *SLC25A39* knockout cells expressing vector or *SLC25A39* cDNA. SLC25A40 data point is highlighted in blue. C. Top 15 scoring genes differentially required for the proliferation of HEK-293T *SLC25A39* knockout cells. D. Relative fold change in cell number of the indicated Jurkat *SLC25A39* knockout cells expressing a vector control or *SLC25A39* cDNA transduced with the indicated sgRNAs. Cells were cultured for 4 days. Cell doublings were normalized to the average of the *SLC25A39* knockout cells expressing *SLC25A39* cDNA (left). Representative bright-field micrographs of Jurkat *SLC25A39* knockout cells transduced with the indicated sgRNA at the end of the cell proliferation assay. Scale bar, 50 μm (right). E. Mitochondrial abundance of GSH in HEK-293T SLC25A39 knockout cells expressing the indicated cDNAs. Data is normalized by CS protein levels. F. Schematic of GSH synthesis in mammals and *S. thermophilus*. Unlike mammals, *S. thermophilus*, express a bifunctional enzyme *GshF*, which possesses both glutamate-cysteine ligase (GCLC) and glutathione synthetase (GS) activities. G. Schematic of engineered GshF construct for mammalian expression and the domains of the protein with GCL and GS function (top). Whole cell GSH abundance (bottom left) and fold change in cell number of HEK-293T cells expressing a vector control and engineered GshF cDNA treated with indicated BSO concentrations for 5 days (bottom right). H. Schematic of engineered GshF construct targeted to mitochondria (mito-GshF) (top). Immunofluorescence analysis of mito-GshF (FLAG, red) and CS (green) in HeLa cells (bottom). Yellow indicates colocalization. Micrographs are representative images. Scale bar, 20 um. I. Mitochondrial abundance of GSH and GSSG in HEK-293T *SLC25A39* knockout cells transduced with the indicated cDNAs. Data is normalized by CS protein levels. J. Schematic showing the restoration of mitochondrial GSH levels by mito-GSHf in *SLC25A39/40* double knockout cells (left). Fold change in cell number (log_2_) of the Jurkat SLC25A39 knockout cells expressing vector or mito-GshF transduced with the indicated sgRNA. Cells were cultured for 4 days (right). K. Fold change in cell number (log_2_) of Jurkat (left) or HEK-293T (right) SLC25A39 knockout cells expressing vector or *SLC25A39* cDNA transduced with the indicated sgRNA. Cells were plated 5 days after transduction and treated for 4 days with the indicated BSO concentrations. L. Synthetic lethal genetic interactions of mtm1 with other genes in *S.cereviase*. **d, e, g, i, j, l,** Bars represent mean ± s.d.; **d, e, g, i, j, l,** *n* = 3 biologically independent samples. Statistical significance in **d**, **e, i, j** was determined by one-way ANOVA followed by Bonferroni post-hoc analysis; **g** and **k** by two-way ANOVA followed by Bonferroni post-hoc analysis.

To formally test whether mitochondrial GSH depletion is necessary for cell proliferation, we would need to restore mitochondrial glutathione levels upon SLC25A39/40 loss. However, modulating glutathione levels specifically in mitochondria is quite challenging. First, GSH synthesis in mammalian cells requires 3 genes (GCLC, GCLM and GS) that encode the GCL holoenzyme and glutathione synthetase^23^. Second, GSH is a non-allosteric feedback inhibitor of γ-glutamate-cysteine ligase, impeding efficient GSH production^24^. Indeed, supplementation of cells with high levels of extracellular glutathione did not influence mitochondrial GSH levels nor did it restore proliferation of SLC25A39/40 double knockout cells (Extended Data Fig 4a,b). These negative results are also consistent with the suggested Km for mitochondrial GSH transport^11^ and the high concentrations of GSH^4^ in the mitochondria.

Several bacterial species including *Streptococcus thermophilus* express a bifunctional enzyme *GshF*, which possesses both glutamate-cysteine ligase and glutathione synthetase activities^25^ (Fig. 3f). *GshF*, unlike the endogenous GCLs, does not display feedback inhibition, allowing efficient GSH accumulation. Building upon these features, we engineered a *GshF* construct that could be utilized to modulate compartmentalized GSH levels (Fig. 3g, Extended Data Fig 4c,d). Confirming the activity of the enzyme in mammalian cells, ectopic expression of a codon-optimized *GshF* cDNA in HEK-293T cells increased cellular GSH levels and conferred resistance to the BSO treatment (Fig. 3g). We next generated a mitochondrially targeted *GshF* (mito-GshF) which faithfully localized to mitochondria and asked whether mito-GshF could functionally complement mitochondrial GSH loss (Fig. 3h, Extended Data Fig. 4e). Remarkably, expression of mito-GshF completely restored the decrease in mitochondrial GSH levels in SLC25A39 knockout cells (Fig. 3i) and rescued the anti-proliferative effects of mitochondrial GSH depletion in SLC25A39/40 double knockout cells (Fig. 3j). Altogether, these results demonstrate that the essential role of SLC25A39/40 is to provide mitochondrial glutathione for cell proliferation.

Given the essential role of mitochondrial GSH import by SLC25A39/40 in proliferation, we predict that mitochondrial GSH depletion should further sensitize cells to a decrease in cellular GSH availability. Indeed, cells lacking SLC25A39/40 die upon inhibition of GSH synthesis by BSO treatment, an effect rescued by SLC25A39 cDNA (Fig. 3k). Similarly, a small-scale sgRNA competition assay in HEK-293T SLC25A39 knockout and control cells infected with a pool of control and SLC25A40 sgRNAs confirmed these results under BSO treatment (Extended Data Fig. 5). Given that SLC25A39/40 is conserved from yeast to higher eukaryotes^26^ (Extended Data Fig. 3b), we next asked whether this association is conserved in lower organisms. Using a yeast interaction network (The Cell Map^27^), we identified GSH1 and GSH2 (yeast glutamylcysteine ligase and glutathione synthetase, respectively) as two of the highest scoring synthetic lethal interactions of mtm1, the *S.cereviase* ortholog for SLC25A39 (p-value=2.78E-28 (GSH1); 7.76E-11(GSH2)) (Fig. 3l). Altogether, these results are consistent with a conserved limiting role of cellular GSH for cell proliferation when mitochondrial GSH import is impaired.

### Loss of mitochondrial GSH import impairs erythropoiesis and the maintenance of iron-sulfur cluster proteins

We next sought to determine the physiological consequences of mitochondrial GSH depletion in mice. In yeast, mtm1 deficiency results in superoxide dismutase defect^28^. In flies, loss of Shawn, the SLC25A39 homolog causes neurodegeneration and synaptic defects^26^. Additionally, SLC25A39 has been shown to co-express with heme synthesis genes and may impact heme metabolism in zebrafish^29^. To determine the organismal function of SLC25A39 in mammals, we generated a full body *Slc25a39-KO* mouse using CRISPR-Cas9 (Extended Data Fig. 6a). Deletion of SLC25A39 was embryonically lethal at day E13.5 and embryos appeared pale, suggesting a severely anemic phenotype (Extended Data Fig. 6b). To more precisely examine the extent to which mitochondrial GSH depletion impacts erythropoiesis, we additionally generated a mouse model in which mitochondrial GSH import is disrupted specifically in the erythroid lineage (conditional *SLC25a39-KO* mice, *ErGFP-cre*; SLC25a39^fl/fl^) (Extended Data Fig. 6c). Mirroring our observation from full body knockouts, these mice also exhibited severe anemia, with a complete lack of Ter119-positive cells and an increased apoptosis in fetal liver cells (Fig. 4a,b; Extended Data Fig. 6d,e). Notably, while we observed a robust reduction in the number of erythroblasts at all differentiation stages, we did not detect a defect in other lineages (Extended Data Fig. 6f,g). In line with these strong phenotypes, SLC25A39 expression is upregulated during red blood cell differentiation *in vivo*, consistent it being the major isoform responsible for GSH import during erythropoiesis (Extended Data Fig. 6h).

**Fig. 4:**
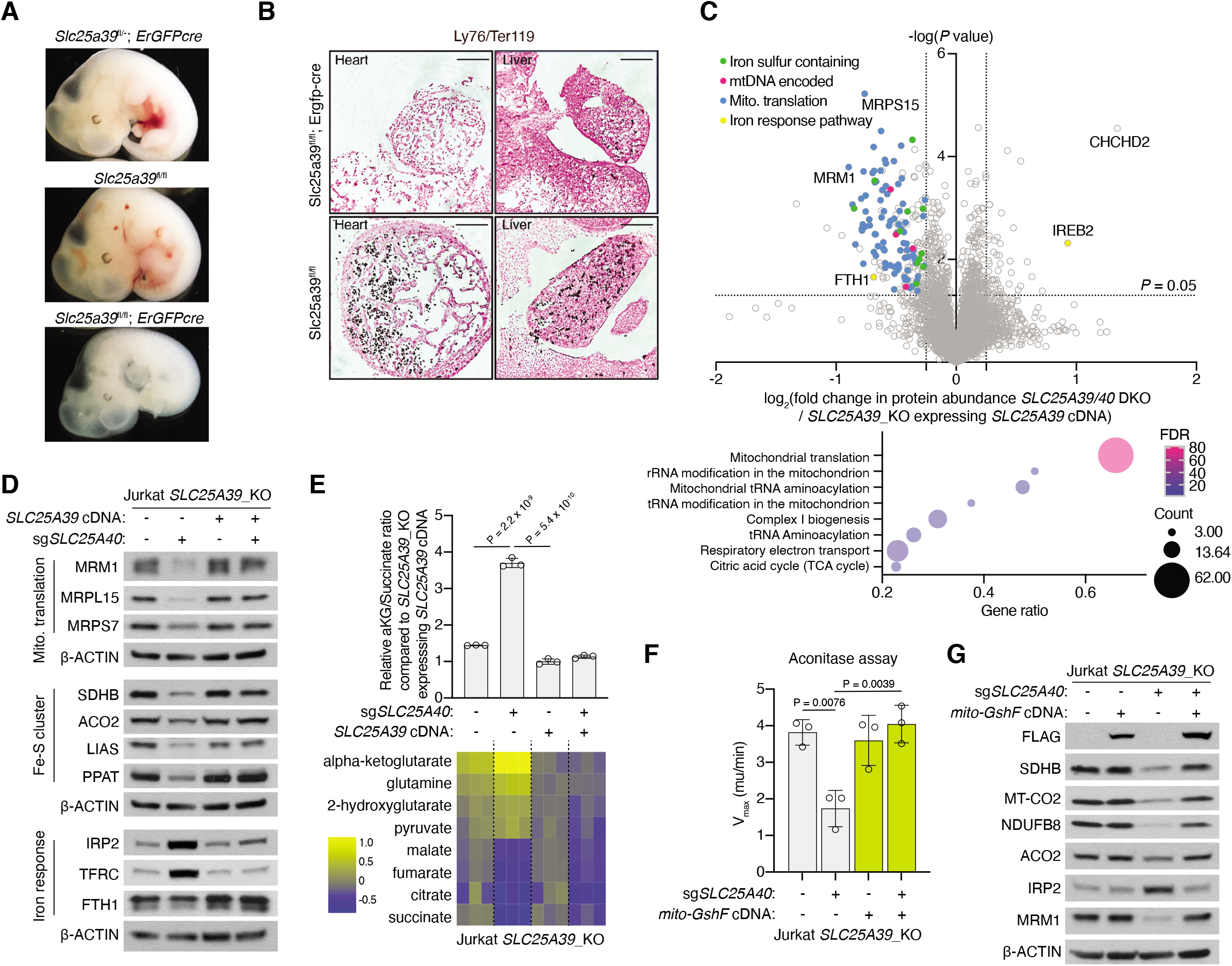
Mitochondrial GSH depletion impairs erythropoiesis and iron-sulfur cluster protein maintenance. A. Representative images for embryos of indicated genotypes at E12.5. Scale bar, 1mm. B. Immunohistochemical staining for developing hearts (left) and livers (right) of E12.5 embryos for indicated genotypes with a Ter119 antibody (black). Nuclei were counterstained with fast red. Scale Bar, 100 μm. C. Volcano plot showing fold change (log_2_) in whole cell protein abundance vs *P* values (-log) for Jurkat *SLC25A39* knockout cells transduced with *sgSLC25A40* compared to those expressing *SLC25A39* cDNA transduced with a control sgRNA (top). Gene ontology analysis of proteins altered by loss of SLC25A39/40 (bottom). D. Immunoblot of indicated proteins in Jurkat *SLC25A39* knockout cells expressing vector or *SLC25A39* cDNA transduced with the indicated sgRNA. Proteins were extracted from the cells 7 days after transduction. β-ACTIN was used as a loading control. E. Metabolomics analysis of Krebs cycle intermediates in Jurkat *SLC25A39* knockout cells expressing vector or *SLC25A39* cDNA transduced with the indicated sgRNA. Metabolites were extracted 5 days after transduction with sgRNAs. Relative ratios of α-ketoglutarate to succinate of each indicated cell line normalized to the average of the SLC25A39 knockout cells expressing *SLC25A39* cDNA (top). Heatmap showing fold change in metabolite levels (log_2_) of the indicated cell lines relative to the average of those in SLC25A39 knockout cells expressing *SLC25A39* cDNA. Values were normalized to the protein concentration of each cell line (bottom). F. Aconitase activity (Vmax) in Jurkat *SLC25A39* knockout cells expressing a vector control or mito-GSHf transduced with the indicated sgRNAs (mean ± SD, n=3). G. Immunoblot of indicated proteins in Jurkat *SLC25A39* knockout cells expressing a vector control or mito-GSHf transduced with the indicated sgRNA. β-ACTIN was used as a loading control. **e, f,** Bars represent mean ± s.d.; **c, e, f,** *n* = 3 biologically independent samples. Statistical significance in **e** and **f** was determined by one-way ANOVA followed by Bonferroni post-hoc analysis.

Despite the essential role of SLC25A39/40 in cell proliferation and the role of SLC25A39 in red blood cell development, how mitochondrial GSH import deficiency impairs cellular functions has never been investigated. To address this mechanistically, we performed an unbiased proteomic analysis on the SLC25A39/40 double knockout Jurkat cells. Our analysis revealed a surprising overlap between proteins downregulated upon BSO treatment and mitochondrial GSH depletion, suggesting that most of the proteomic changes that occur upon cellular GSH depletion are likely due to mitochondrial effects (Fig. 4c, Extended Data Fig. 7a). Indeed, gene ontology analysis for downregulated proteins in SLC25A39/40 double knockout cells revealed a strong enrichment for mitochondrial translation and iron-sulfur cluster containing proteins (Fig. 4c, Extended Data Fig. 7b). Additionally, we observed an induction of iron regulatory protein 2 (IRP2), a major component of the iron response pathway which is activated upon iron-sulfur cluster synthesis dysfunction^30^ (Fig. 4c). We confirmed these proteomic results in individual immunoblotting experiments using Jurkat and HEK-293T cells (Fig. 4d, Extended Data Fig. 7c). Importantly, these effects could be recapitulated in cysteine desulfurase (NFS1) knockdown cells with impaired iron-sulfur cluster synthesis, indicating that phenotypes observed in SLC25A39/40 knockout cells are likely a consequence of an iron-sulfur cluster deficiency (Extended Data Fig. 7d). Consistent with an iron-sulfur cluster defect, SLC25A39/40 knock out cells displayed reduced activity of the enzyme aconitase, in which iron sulfur clusters function in both substrate binding and catalysis (Fig 4f). Additionally, SLC25A39/40 double knockout cells exhibit reduction in the protein levels of lipoic acid synthase (LIAS), another iron-sulfur cluster containing protein involved in the synthesis of lipoate^31^. Lipoate is an essential lipid modification that is required for catalysis by the α-ketoglutarate dehydrogenase (OGDH) and pyruvate dehydrogenase (PDH) complexes. Consistent with a inhibition at the PDH and OGDH, SLC25A39/40 double knockout cells accumulated α-ketoglutarate (aKG) and displayed elevations in the aKG/succinate and pyruvate/citrate ratios (Fig 4e, Extended Data Fig. 7e). Strikingly, simply increasing mitochondrial GSH levels by expressing mito-GSHf in SLC25A39/40 double knockout cells completely restored the reduced levels of mitochondrial translation machinery proteins, iron-sulfur containing proteins, and aconitase activity (Fig 4f,g). Notably, addition of hemin or pyruvate, a source for NAD+ did not impact these growth phenotypes, suggesting that amelioration of heme defect or respiration inhibition is not sufficient to overcome anti-proliferative effects of mitochondrial GSH depletion (Extended Data Fig. 7f). Therefore, we conclude that mitochondrial GSH depletion impairs the activity and stability of iron sulfur cluster containing proteins.

## Discussion

Our work identifies SLC25A39 and its paralogue SLC25A40 as mitochondrial carriers required for GSH import. A previous report suggested that mitochondrial dicarboxylate and 2-oxoglutarate carriers may additionally transport glutathione^32^, a finding later disproved^33^. Similarly, expression of these transporters was unable to restore the decrease in glutathione levels in our assays (Extended Data Fig. 8a). Given the essential role of SLC25A39 in mitochondrial glutathione transport, our results provide a mechanistic explanation for how SLC25A39 loss causes mitochondrial defects, neurodegeneration and anemia in yeast, flies and mice, respectively. Mutations that impair glutathione synthesis are associated with rare forms of anemia in humans^34^. Interestingly, a *SLC25A39*-associated human phenome analysis using 4,000 traits in the UK Biobank (n=361,194 subjects) revealed reticulocyte count and hemoglobin concentration as the top significant phenotypes associated with genetically regulated *SLC25A39* expression (Extended Data Fig. 8b). Additionally, an SLC25A40 variant is associated with hyperlipidemia^35^. Additionally, SLC25A39 expression is induced in brown fat in response to cold^36^, indicating that it may also be regulated by physiological cues. These diverse phenotypes are consistent with the differential expression patterns of SLC25A39 and SLC25A40 across tissues. Future work is required to dissect these precise physiological roles and regulation of these transporters in organismal studies.

Mitochondrial GSH import by SLC25A39/40 is necessary for maintaining steady state levels of iron-sulfur clusters. The critical need for mitochondrial GSH in iron-sulfur clusters is consistent with its role as a cofactor for mitochondrial glutaredoxins (Grxs), which are conserved oxidoreductases for iron-sulfur cluster biogenesis. Finally, SLC25A39 is one of the few mitochondrial carriers that is feedback-regulated at the protein level by a downstream metabolite. Given that this regulation is independent of NRF2, our results suggest the presence of a potential signaling pathway regulated by GSH availability. Identification of molecular factors regulating SLC25A39 protein levels and mitochondrial GSH import will provide insights into how compartmentalized GSH levels may be sensed in mammalian cells. Future structural and biochemical studies will allow for precise delineation of how mitochondrial GSH transport is controlled by upstream cues.

## Acknowledgements

We thank all members of the Birsoy lab for helpful suggestions. K.B. was supported by NCI (DP2 OD024174-01) and Mark Foundation Emerging Leader Award; and is a Searle and Pew-Stewart Scholar.

## Extended Figure Legends

**Extended Data Fig. 1: SLC25A39 protein levels are directly regulated by cellular GSH availability.**

A. Gene enrichment analysis (Gorilla) for proteomics data from the immune isolated mitochondria of HeLa cells.

B. Immunoblot of SLC25A39, SLC25A12 and CS in HeLa cells treated with the indicated doses of BSO for 24 hours. CS was used as the loading control.

C. Relative abundance of *SLC25A39* and *GCLM* transcripts in HEK-293T cells treated with BSO (1 mM) for the indicated days using quantitative reverse transcription PCR (RT-qPCR), normalized to transcripts of *B-ACTIN*. Error bars represent mean ± s.d.; *n* = 3 biologically independent samples. Statistical significance was determined by One-way ANOVA followed by Bonferroni post-hoc analysis.

D. Immunoblot of indicated proteins in HEK-293T *SLC25A39* knockout cells expressing a vector control or SLC25A39 cDNA treated with BSO (1 mM) for the indicated times. GAPDH was used as the loading control.

E. Immunoblot of indicated proteins in HeLa cells treated with indicated doses of H_2_O_2_ for 24 hours. CS and GAPDH were used as loading controls.

F. Immunoblot of indicated proteins in HeLa cells treated with Erastin (5 μM) or indicated doses of KI696 (NRF2 activator) for 24 hours. GAPDH were used as loading controls.

G. Immunoblot of indicated proteins in HeLa cells treated with Erastin (5 μM) or indicated doses of RSL-3, an inhibitor of glutathione peroxidase 4 (GPX4) for 24 hours. CS and GAPDH were used as loading controls.

H. Immunoblot of indicated proteins in HeLa cells treated with Erastin (5 μM) and co-treated with either GSH (10 mM), GSHee (10 mM), Trolox (50 uM) or Ferrostatin-1 (5 uM) for 48 hours. CS was used as a loading control.

**Extended Data Fig. 2: Mitochondrial GSH import is mediated by SLC25A39.**

A. Volcano plots showing the fold change in mitochondrial metabolite abundance (log_2_) vs *P* values (-log) from K562 SLC25A39 knockout cell lines expressing a vector control or *SLC25A39* cDNA. Red data points highlight GSH and GSSG. The dotted line represents *P*=0.05.

B. Immunoblot of indicated proteins in whole-cell lysates and mitochondria isolated from HEK-293T (top) and HeLa (bottom) SLC25A39 knockout cells expressing a vector control or *SLC25A39* cDNA.

C. Mitochondrial GSH and GSSG abundance (top and middle panels) and total intracellular GSH levels in parental HEK-293T cells and *SLC25A39* knockout cells expressing a vector control or *SLC25A39* cDNA. Data is normalized by citrate synthase (CS) protein levels.

D. Uptake of indicated concentrations of [^13^C_2_,^15^N]-GSH into mitochondria isolated from parental HEK-293T mitotag cells for 10 minutes. Data is normalized to the NAD+ abundance.

E. Uptake of indicated concentrations of [^13^C_2_,^15^N]-GSH or GSH into mitochondria isolated from parental HEK-293T mitotag cells for 10 minutes. Data is normalized to the NAD+ abundance.

**c**, **d**, **e**, Bars represent mean ± s.d.; **a, c**, **d, e,** *n* = 3 biologically independent samples. Statistical significance in **c** and **d** was determined by One-way ANOVA followed by Bonferroni post-hoc analysis; **e** by two-tailed unpaired *t*-test.

**Extended Data Fig. 3: SLC25A40, the mammalian paralog for SLC25A39 can replace SLC25A39 loss.**

A. Individual sgRNA scores of SLC25A40 and TXNRD2 from the CRISPR screens in indicated cell lines from Fig. 3b.

B. Phylogenetic tree of SLC25A39 homologs across model organisms.

C. Top ranked (z-scores) co-evolved genes with SLC25A40 across species. (phylogene)

D. Relative conservation of SLC25A40 and SLC25A39 in different species compared to their human homologs.

E. Relative fold change in cell number of the indicated HEK-293T SLC25A39 knockout cells expressing vector or *SLC25A39* cDNA transduced with the indicated sgRNAs. Cells were plated 5 days after transduction and cultured for 4 days (mean ± SD, n=3). Cell doublings were normalized to the average of the SLC25A39 knockout cells expressing *SLC25A39* cDNA. Statistical significance was determined by One-way ANOVA followed by Bonferroni post-hoc analysis.

F. Volcano plots of the fold change in mitochondrial metabolite abundance (log_2_) vs *P* values (-log) from HEK-293T SLC25A39 knockout cell line expressing a vector control or *SLC25A40* cDNA. Red data points highlight GSH and GSSG. Red data points highlight GSH and GSSG. The dotted line represents *P*=0.05.

**Extended Data Fig. 4: Expression of *S.thermophilus* GshF can modulate cellular GSH levels in mammalian cells.**

A. Mitochondrial abundance of GSH in HEK-293T *SLC25A39* knockout cells expressing a vector control or *SLC25A39* cDNA treated with GSH (5 mM) or GSHee (5 mM) for 24 hours. Data is normalized by mitochondrial NAD+.

B. Relative cell number of indicated SLC25A39 knockout Jurkat cells transduced with indicated sgRNAs. Cells were plated 5 days after transduction and cultured for 4 days with or without GSHee (10 mM). Cell numbers were normalized to the average of the SLC25A39 knockout cells in each treatment condition.

C. Immunofluorescence analysis of GshF (FLAG, red) and CS (green) in HeLa cells (bottom). Micrographs are representative images. Scale bar, 20 μm.

D. Immunoblots of indicated proteins in HEK-293T cells expressing a vector control or GshF cDNA. GAPDH was used as the loading control.

E. Immunoblot analysis of indicated proteins in whole-cell lysates and mitochondria isolated from SLC25A39 knockout HEK-293T cells expressing a vector control, mito-GshF or SLC25A39 cDNA.

**a, b,** Bars represent mean ± s.d.; **a, b,** *n* = 3 biologically independent samples. Statistical significance in **a** and **b** was determined by One-way ANOVA followed by Bonferroni post-hoc analysis.

**Extended Data Fig. 5: Mitochondrial GSH depletion sensitizes cells to BSO treatment.**

Scheme of the *in vitro* sgRNA competition assay performed in HEK-293T *SLC25A39* knockout cells expressing a vector control or *SLC25A39* cDNA transduced with a pool of 5 control sgRNAs (sgControl, gray) and sgRNAs targeting TXNRD2 (pink) and SLC25A40 (green) (left). Differential guide scores in the indicated cell lines upon treatment with 20 μM BSO (right). Bars represent mean ± s.d.; Statistical significance was determined by two-tailed unpaired *t*-test.

**Extended Data Fig. 6: Slc25a39 is essential for embryonic development and red cell differentiation *in vivo*.**

A. Targeting scheme for Slc25a39 knockout mice. Bar, 1mm.

B. Gross appearance of E12.5 embryos of the indicated genotypes. Genotyping of Slc25a39 E12.5 embryos from heterozygous mating. PCR of wild type allele and targeted allele result in ~1500bp and ~800bp bands, respectively. The number of viable pups with indicated genotypes is shown (right).

C. Targeting scheme for Slc25a39 conditional knockout (Slc25a39^fl/fl^) mice in which two loxp sites were inserted in the indicated intron regions. The resultant Slc25a39^fl/fl^ mice were mated with erythroid-lineage specific Cre-recombinase (Ergfp-cre) mice to generate an erythroid-specific conditional knockout mice.

D. The number of viable pups with indicated genotypes is shown resulting from the indicated mating.

E. Representative images of H&E staining of fetal liver cells from E12.5 Slc25a39^fl/fl^ and Ergfpcre^+/-^ Slc25a39^fl/fl^ embryos. The bottom images are the boxed area of the top images. Arrows show that many hematopoietic cells in the fetal liver display nuclear fragmentation and cellular shrinkage. Bars, 50μm (top panel), 12.5μm (bottom panel).

F. Quantification of erythroblast populations at different differentiation stages using surface markers (bottom): Ter119 and CD44, and FSC (size). Data are shown as the percentage of cells of indicated populations among total fetal liver cells. Gating strategy (top). Bars represent mean ± s.d.; *n* = 3, Ergfp-cre^+/-^ Slc25a39^fl/fl^; 4 Slc25a39^fl/fl^ embryos. Statistical significance was determined by One-way ANOVA followed by Bonferroni post-hoc analysis.

G. Profiling of cells of myeloid lineage (top), progenitor cells (middle) and of lymphoid lineage (bottom). Data were presented as the population among total fetal liver cells. Bars represent mean ± s.d.; *n* = 12 Slc25a39^fl/fl^ or Slc25a39^fl/-^, 7 Ergfp-cre^+/-^ Slc25a39^fl/-^ and 4 Ergfp-cre^+/-^ Slc25a39^fl/fl^ E12 embryos. Statistical significance was determined by One-way ANOVA followed by Bonferroni post-hoc analysis.

H. Relative RPKM values of SLC25A39, FTL and HBB genes in human erythroblasts at differing stages of terminal differentiation.

**Extended Data Fig. 7: Loss of SLC25A39/40 decreases the steady state levels of iron-sulfur containing proteins and phenocopies iron-sulfur cluster deficiency.**

A. Comparison of proteomics data from BSO treated HeLa cells and *SLC25A39/40* double knockout Jurkat cells. The criteria are indicated on the venn diagram.

B. Gene ontology analysis of significantly downregulated genes in *SLC25A39/40* double knockout Jurkat cells.

C. Immunoblot of indicated proteins in the indicated HEK-293T SLC25A39 knockout cells expressing vector or *SLC25A39* cDNA transduced with the indicated sgRNA. Proteins were extracted from the cells 7 days after transduction. β-ACTIN was used as a loading control.

D. Immunoblot of indicated proteins in indicated HEK-293T cells transduced with shRNA targeting either GFP as a control or NFS1 (two different shRNAs). Proteins were extracted from cells 6 days after transduction. GAPDH was used as a loading control.

E. Schematic showing how the iron-sulfur clusters-containing protein LIAS enables OGDH and PDH activity by transferring lipoic acid as a cofactor to their enzyme complexes (top left). Relative ratios of pyruvate to citrate metabolite levels of each indicated cell line from the metabolomics analysis in Figure 4E. Ratios were normalized to the average of the *SLC25A39* knockout cells expressing *SLC25A39* cDNA (bottom left). Immunoblot with antibody against lipoic acid in the indicated Jurkat *SLC25A39* knockout cells expressing vector or *SLC25A39* cDNA transduced with the indicated sgRNA. Arrows indicate E2 complexes of PDH and OGDH containing lipoic acid (right).

F. Representative bright-field micrographs of Jurkat SLC25A39 knockout cells transduced with the indicated sgRNA at the end of the cell proliferation assay. Scale bar, 50 μm.

G. Relative cell number of the indicated Jurkat *SLC25A39* knockout cells expressing a vector control or *SLC25A39* cDNA transduced with the indicated sgRNA. Cells were plated 5 days after transduction and cultured for 4 days with or without hemin (1 μM) and pyruvate (1 mM). Cell numbers were normalized to the average of the SLC25A39 knockout cells in each treatment condition.

**e**, **g**, Bars represent mean ± s.d.; **e**, **g**, *n* = 3 biologically independent samples. Statistical significance in **e** and **g** was determined by One-way ANOVA followed by Bonferroni post-hoc analysis.

**Extended Data Fig. 8: Expression of DIC/OGC does not complement the decrease of mitochondrial GSH availability in SLC25A39 knockout cells.**

A. Uptake of [^13^C_2_,^15^N]-GSH into mitochondria isolated from HEK-293T *SLC25A39* knockout cells expressing a vector control, SLC25A39 cDNA, SLC25A10 (DIC, mitochondrial dicarboxylate carrier) or SLC25A11 (OGC, mitochondrial oxoglutarate carrier) cDNA at 5 seconds and 10 minutes. Bars represent mean ± s.d.; *n* = 3 biologically independent samples. Statistical significance was determined by Two-way ANOVA followed by Bonferroni post-hoc analysis.

B. PrediXcan-based TWAS design in UK biobank and summary graph of SLC25A39-associated phenotypes. Bonferroni-adjusted p-values are reported.

## MATERIALS AND METHODS

### Cell lines and reagents

Human cells lines HeLa, Jurkat and HEK-293T were purchased from the ATCC. Cell lines were verified to be free of mycoplasma contamination and the identities of all were authenticated by STR profiling. Jurkat cells were maintained in RPMI media (Gibco) containing 2 mM glutamine, 10% fetal bovine serum, 1% penicillin and streptomycin. HeLa and HEK-293T cells were maintained in either DMEM media (Gibco) containing 4.5g/L glucose, 110mg/L pyruvate, 4mM glutamine, 10% fetal bovine serum, 1% penicillin and streptomycin or in RPMI media (Gibco) containing 2 mM glutamine, 10% fetal bovine serum, 1% penicillin and streptomycin.

Antibodies against beta-Actin (GTX109639), GAPDH (GTX627408) and PPAT (GTX102725) were obtained from GeneTex; lipoic acid antibody (437695) from EMD Millipore; SLC25A39 (14963-1-AP), ALAS1 (16200-1-AP), GCLC (12601-1-AP), HMOX1 (10701-1-AP), MRM1 (16392-1-AP), MRPL15 (18339-1-AP), MRPS7 (26828-1-AP), FDX1 (12592-1-AP), FECH (14466-1-AP) and LIAS (11577-1-AP) antibodies from Proteintech; anti-FLAG M2 (F1804) from Sigma; total OXPHOS antibody cocktail (ab110411) and SDHB (ab14714) antibody from Abcam; NFS1 (sc-365308) from Santa Cruz Biotechnology; SLC25A12 (64169S), calreticulin (12238P), RCAS1 (12290S), LAMP1 (9091P), CS (14309S), ACO2 (6571S), IRP2 (37135S), TFRC (13208S), FTH1 (3998S), cleaved PARP (9546S), SLC7A11 (12691S) and RPS6 (2217S) antibodies from CST; and ATP5A (43-9800) from ThermoFisher Scientific.

[13C2,15N]-GSH was purchased from Cambridge Isotope Laboratory (CNLM-6245-50), L(-)-Glutathione (reduced form) (MP Biomedicals, 0210181401). anti-HA magnetic beads (Thermo Scientific Pierce 88837). DAPI (D1306) was purchased from ThermoFisher Scientific; sodium pyruvate (P2256), polybrene (H9268), puromycin (P8833), Glutathione reduced ethyl ester (G1404), Hemin (H9039), BSO (B2515) from Sigma; blasticidin from Invivogen (ant-bl-1); erastin (5449) from Tocris Bioscience.

### Generation of knockout, knockdown and cDNA overexpression cell lines

sgRNAs (oligonucleotide sequences are indicated below) were cloned into lentiCRISPR-v2 (Addgene) (for *GCLC*)or into lentiCRISPR-v1 (Addgene) (for *SLC25A39*) linearized with BsmBI by T4 ligase (NEB). sgRNA expressing vector along with lentiviral packaging vectors Delta-VPR and CMV VSV-G were transfected into HEK-293T cells using the XTremeGene 9 transfection reagent (Roche). Similarly, for overexpression cell lines, gBlocks (IDT) containing the cDNA of interest were cloned into pMXS-IRES-Blast linearized with BamHI and NotI by Gibson Assembly (NEB). cDNA vectors along with retroviral packaging vectors gag-pol and CMV VSV-G were transfected into HEK-293T cells. The virus-containing supernatant was collected 48 hrs after transfection and passed through a 0.22 μm filter to eliminate cells. Target cells in 6-well tissue culture plates were infected in media containing 8 μg/mL of polybrene and a spin infection was performed by centrifugation at 2,200 rpm for 1 hour. Post-infection, virus was removed and cells were selected with puromycin (lentiCRISPR-v2), blasticidin (pMXS_IRES_Blast) or by flow sorting of top 1% GFP+ cells (lentiCRISPR-v1(GFP)). Clones were validated for loss of the relevant protein via immunoblotting. As no valid antibody is available, sgSLC25A40 transduced cells were validated through ICE analysis (Synthego) of gDNA Sanger sequencing. Unless otherwise stated, double knockout of SLC25A39_KO cells with sgSLC25A40 was lethal and each experiment was done with freshly transduced cells after the indicated number of days. For NFS1 knock down, the following shRNAs were used: shNFS1_1, TRCN0000229753, B4: shNFS1_2, TRCN0000229755, B6 (Addgene).

#### Oligo Sequences

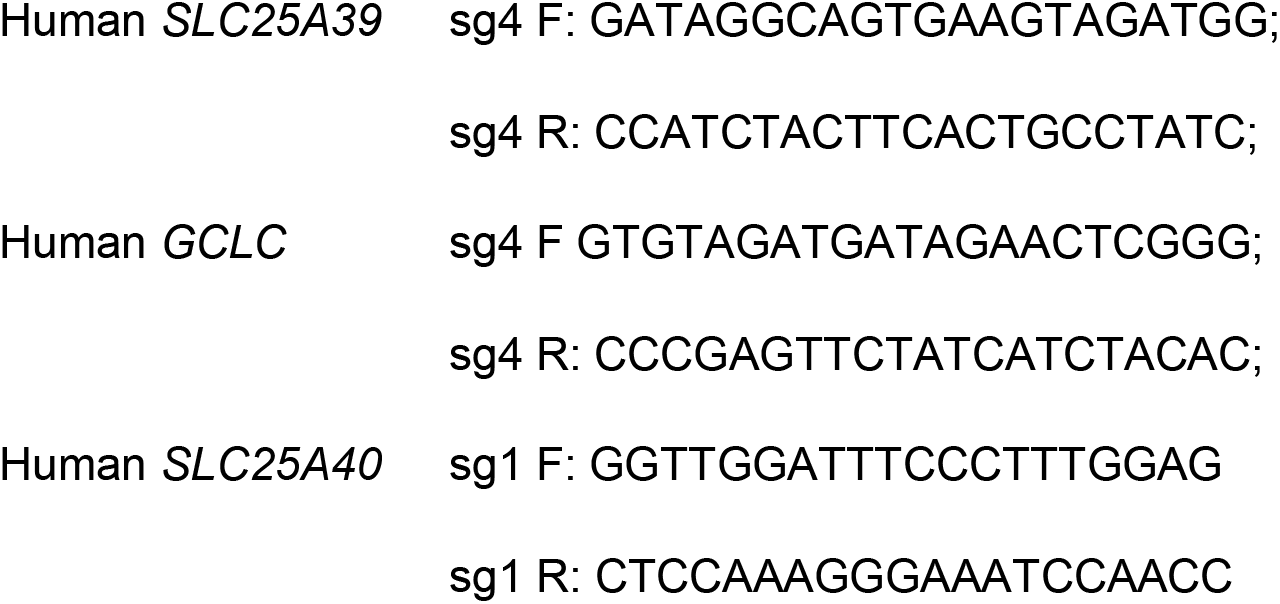

#### Codon-optimized *SLC25A39* sequence

ATGGCAGACCAGGACCCCGCGGGCATCTCACCTCTCCAGCAGATGGTCGCATCTGGAACAGGGGCAGTCGTCACAAGTTTGTTCATGACCCCACTTGATGTAGTGAAAGTCCGGCTTCAATCACAACGCCCTAGCATGGCCAGCGAGCTGATGCCGAGCTCCAGGCTCTGGTCACTTTCTTATACGAAGCTTCCCTCTTCTCTCCAGTCTACGGGTAAATGTTTGCTTTATTGTAACGGCGTACTCGAACCTCTGTATTTGTGTCCAAATGGAGCACGCTGCGCCACGTGGTTTCAGGACCCAACTCGATTTACCGGCACAATGGACGCATTTGTCAAGATAGTAAGACACGAGGGTACAAGAACGCTTTGGAGCGGCCTCCCTGCTACGTTGGTGATGACGGTTCCCGCAACGGCCATATACTTTACAGCCTACGACCAGCTGAAGGCCTTTCTGTGTGGTAGGGCACTTACCTCAGACCTTTACGCTCCAATGGTCGCAGGGGCCCTTGCAAGACTTGGTACGGTCACTGTAATAAGTCCGCTCGAACTCATGAGGACAAAACTCCAAGCTCAGCACGTGAGCTACCGGGAACTGGGGGCTTGTGTACGCACAGCGGTCGCGCAAGGCGGCTGGAGGAGTCTGTGGCTGGGTTGGGGGCCCACGGCCCTCCGGGACGTACCGTTTTCTGCGCTTTATTGGTTTAACTACGAGCTTGTGAAATCTTGGCTCAATGGATTCCGGCCGAAAGACCAGACCTCCGTTGGAATGTCTTTCGTCGCCGGGGGCATTTCCGGCACGGTGGCCGCCGTGCTGACCTTGCCATTCGACGTTGTTAAGACCCAGCGACAGGTCGCTTTGGGGGCAATGGAGGCCGTGCGGGTGAACCCACTCCACGTTGACAGTACATGGTTGCTGCTCCGCCGCATCCGGGCCGAAAGCGGAACTAAAGGTCTGTTTGCTGGATTTCTTCCGCGAATCATTAAGGCTGCGCCATCTTGTGCAATCATGATCTCTACATACGAGTTTGGAAAATCCTTCTTTCAGAGGCTTAATCAGGACAGACTGCTCGGAGGGTAA

#### *SLC25A39* D226A

ATGGCAGACCAGGACCCCGCGGGCATCTCACCTCTCCAGCAGATGGTCGCATCTGGAACAGGGGCAGTCGTCACAAGTTTGTTCATGACCCCACTTGATGTAGTGAAAGTCCGGCTTCAATCACAACGCCCTAGCATGGCCAGCGAGCTGATGCCGAGCTCCAGGCTCTGGTCACTTTCTTATACGAAGCTTCCCTCTTCTCTCCAGTCTACGGGTAAATGTTTGCTTTATTGTAACGGCGTACTCGAACCTCTGTATTTGTGTCCAAATGGAGCACGCTGCGCCACGTGGTTTCAGGACCCAACTCGATTTACCGGCACAATGGACGCATTTGTCAAGATAGTAAGACACGAGGGTACAAGAACGCTTTGGAGCGGCCTCCCTGCTACGTTGGTGATGACGGTTCCCGCAACGGCCATATACTTTACAGCCTACGACCAGCTGAAGGCCTTTCTGTGTGGTAGGGCACTTACCTCAGACCTTTACGCTCCAATGGTCGCAGGGGCCCTTGCAAGACTTGGTACGGTCACTGTAATAAGTCCGCTCGAACTCATGAGGACAAAACTCCAAGCTCAGCACGTGAGCTACCGGGAACTGGGGGCTTGTGTACGCACAGCGGTCGCGCAAGGCGGCTGGAGGAGTCTGTGGCTGGGTTGGGGGCCCACGGCCCTCCGGGCAGTACCGTTTTCTGCGCTTTATTGGTTTAACTACGAGCTTGTGAAATCTTGGCTCAATGGATTCCGGCCGAAAGACCAGACCTCCGTTGGAATGTCTTTCGTCGCCGGGGGCATTTCCGGCACGGTGGCCGCCGTGCTGACCTTGCCATTCGACGTTGTTAAGACCCAGCGACAGGTCGCTTTGGGGGCAATGGAGGCCGTGCGGGTGAACCCACTCCACGTTGACAGTACATGGTTGCTGCTCCGCCGCATCCGGGCCGAAAGCGGAACTAAAGGTCTGTTTGCTGGATTTCTTCCGCGAATCATTAAGGCTGCGCCATCTTGTGCAATCATGATCTCTACATACGAGTTTGGAAAATCCTTCTTTCAGAGGCTTAATCAGGACAGACTGCTCGGAGGGTAA

#### *SLC25A39* K329A

gccaccATGGCAGACCAGGACCCCGCGGGCATCTCACCTCTCCAGCAGATGGTCGCATCTGGAACAGGGGCAGTCGTCACAAGTTTGTTCATGACCCCACTTGATGTAGTGAAAGTCCGGCTTCAATCACAACGCCCTAGCATGGCCAGCGAGCTGATGCCGAGCTCCAGGCTCTGGTCACTTTCTTATACGAAGCTTCCCTCTTCTCTCCAGTCTACGGGTAAATGTTTGCTTTATTGTAACGGCGTACTCGAACCTCTGTATTTGTGTCCAAATGGAGCACGCTGCGCCACGTGGTTTCAGGACCCAACTCGATTTACCGGCACAATGGACGCATTTGTCAAGATAGTAAGACACGAGGGTACAAGAACGCTTTGGAGCGGCCTCCCTGCTACGTTGGTGATGACGGTTCCCGCAACGGCCATATACTTTACAGCCTACGACCAGCTGAAGGCCTTTCTGTGTGGTAGGGCACTTACCTCAGACCTTTACGCTCCAATGGTCGCAGGGGCCCTTGCAAGACTTGGTACGGTCACTGTAATAAGTCCGCTCGAACTCATGAGGACAAAACTCCAAGCTCAGCACGTGAGCTACCGGGAACTGGGGGCTTGTGTACGCACAGCGGTCGCGCAAGGCGGCTGGAGGAGTCTGTGGCTGGGTTGGGGGCCCACGGCCCTCCGGGACGTACCGTTTTCTGCGCTTTATTGGTTTAACTACGAGCTTGTGAAATCTTGGCTCAATGGATTCCGGCCGAAAGACCAGACCTCCGTTGGAATGTCTTTCGTCGCCGGGGGCATTTCCGGCACGGTGGCCGCCGTGCTGACCTTGCCATTCGACGTTGTTAAGACCCAGCGACAGGTCGCTTTGGGGGCAATGGAGGCCGTGCGGGTGAACCCACTCCACGTTGACAGTACATGGTTGCTGCTCCGCCGCATCCGGGCCGAAAGCGGAACTAAAGGTCTGTTTGCTGGATTTCTTCCGCGAATCATTGCAGCTGCGCCATCTTGTGCAATCATGATCTCTACATACGAGTTTGGAAAATCCTTCTTTCAGAGGCTTAATCAGGACAGACTGCTCGGAGGGTAA

These gene fragments were purchased from Twist Biosciences, and directly cloned into linearized pMXS-IRES-Blast (Cell Biolabs, RTV-016).

### GshF Construct

Below are *H. sapiens* codon optimized sequences of *S. thermophilus* GshF with added mitochondrial targeting sequences (MTS) and FLAG-tags. To ensure mitochondrial localization of the mito-GshF construct, a tandem MTS consisting of the *L. brevis* COXIV MTS followed by the *Homo sapiens* ACOI MTS was added to the N-terminus of the GshF sequence, along with an N-terminal FLAG-tag. For clarity, corresponding regions of each sequence are highlighted as follows: FLAG tag, MTS, (GGS)_3_ linker, Stop codo, Gibson overhangs.

#### Codon optimized GshF

GCCGGATCTAGCTAGTTAATTAAGccaccatgACCCTTAATCAGCTTCTCCAGAAGTTGGAGGCGACTTCCCCCATTCTCCAGGCGAACTTCGGGATAGAAAGGGAGTCATTGAGGGTTGACCGCCAGGGTCAGCTGGTCCACACACCGCACCCCTCATGTCTGGGAGCCCGCAGTTTTCATCCTTACATACAAACCGACTTTTGTGAATTCCAAATGGAACTGATTACACCAGTAGCCAAAAGTACGACGGAGGCCCGACGCTTTCTTGGCGCGATAACTGATGTAGCAGGACGAAGCATTGCAACTGACGAGGTGCTGTGGCCATTGAGTATGCCACCACGACTTAAAGCCGAGGAAATTCAAGTAGCGCAACTCGAGAACGACTTCGAAAGACATTATCGGAACTACTTGGCAGAGAAGTACGGCACCAAATTGCAGGCGATTAGTGGAATTCATTACAATATGGAACTTGGGAAGGACTTGGTTGAGGCGCTTTTTCAAGAGTCAGATCAGACTGACATGATCGCATTTAAAAACGCTCTGTATCTCAAGCTCGCCCAGAACTATTTGAGGTATCGGTGGGTCATTACTTACCTGTTTGGAGCAAGTCCCATTGCAGAACAAGGATTCTTTGACCAAGAAGTGCCGGAGCCTATGCGCTCTTTCCGCAACTCCGACCACGGCTACGTTAACAAGGAAGAGATACAGGTAAGCTTTGTATCCCTTGAAGACTATGTCTCCGCGATCGAGACCTACATCGAGCAGGGTGACCTTATAGCCGAGAAAGAGTTTTACTCAGCCGTGCGCTTTAGAGGACAAAAAGTCAATCGCTCCTTCCTTGATAAGGGTATAACTTATCTGGAGTTCAGAAACTTTGACTTGAACCCATTTGAGAGAATAGGCATCAGTCAAACCACTATGGATACCGTTCACTTGCTCATACTGGCCTTTCTCTGGTTGGATAGTCCGGAAAACGTGGACCAGGCCCTTGCCCAGGGACACGCGCTTAACGAGAAAATAGCCCTCTCCCATCCATTGGAGCCCCTCCCTTCAGAGGCCAAGACACAGGACATCGTGACCGCACTCGACCAGCTCGTACAGCACTTCGGATTGGGAGATTACCACCAGGATCTCGTGAAACAAGTGAAAGCGGCGTTTGCTGATCCGAATCAAACCCTGTCAGCTCAACTTCTTCCTTATATTAAGGACAAGTCACTCGCAGAATTCGCTCTCAATAAGGCACTCGCATATCATGACTATGACTGGACCGCTCACTACGCCCTTAAAGGTTACGAAGAAATGGAGCTCAGTACGCAGATGCTGCTCTTTGATGCTATCCAGAAAGGAATACACTTCGAGATACTCGATGAGCAAGATC AGTTCTTGAAGCTGTGGCATCAAGATCACGTAGAATATGTTAAAAACGGTAATATGACCAGCAAGGATAACTATGTAGTACCTCTCGCAATGGCCAACAAGACTGTTACTAAGAAAATTCTTGCTGACGCTGGGTTCCCTGTTCCGTCCGGGGACGAATTTACTAGCTTGGAGGAGGGACTGGCCTACTACCCGCTTATTAAAGATAAGCAAATTGTAGTAAAGCCAAAGAGCACGAATTTCGGCTTGGGTATCAGCATCTTCCAAGAACCCGCCAGTCTCGACAATTATCAAAAAGCATTGGAAATAGCATTTGCGGAGGACACTAGTGTGCTCGTCGAAGAATTCATTCCAGGCACGGAATACCGATTCTTCATTTTGGACGGACGCTGTGAGGCAGTCCTTTTGAGGGTAGCTGCCAATGTAATAGGGGACGGGAAACACACAATCAGAGAGTTGGTAGCGCAGAAAAACGCAAATCCCCTGCGCGGTAGGGATCATAGATCACCCTTGGAAATCATAGAGCTTGGGGACATAGAGCAACTCATGCTGGCACAGCAGGGTTACACTCCAGATGACATCCTGCCAGAGGGTAAGAAAGTGAATTTGAGGCGGAACAGCAATATTAGTACTGGGGGAGACTCCATAGACGTCACAGAAACAATGGATAGCTCTTATCAAGAACTTGCAGCAGCGATGGCTACCAGTATGGGGGCATGGGCCTGTGGAGTTGATCTGATTATACCCGACGAAACGCAGATTGCCACAAAGGAAAATCCACATTGCACGTGTATTGAACTTAACTTCAACCCCTCCATGTACATGCATACATACTGCGCTGAGGGGCCGGGGCAGGCAATTACAACCAAAATACTCGACAAACTCTTCCCGGAGATCGTGGCCGGACAAACTGGAGGAAGCGGAGGAAGCGGAGGAAGCGATTACAAGGATGACGATGACAAGTAAGCTACGTAAATTCC GCCC

### Mito-GSHf

GCCGGATCTAGCTAGTTAATTAAGccaccATGCTCGCTACAAGGGTCTTTAGCCTCGTCGGAAAGAGAGCTATCAGCACCTCCGTCTGCGTGAGAGCTCATatggcgccctacagcctactggtgactcggctgcagaaagctctgggtgtgcggcagtaccatgtggcctcagtcctgtgcGATTACAAGGATGACGATGACAAGGGAGGAAGCGGAGGAAGCGGAGGAAGCACCCTTAATCAGCTTCTCCAGAAGTTGGAGGCGACTTCCCCCATTCTCCAGGCGAACTTCGGGATAGAAAGGGAGTCATTGAGGGTTGACCGCCAGGGTCAGCTGGTCCACACACCGCACCCCTCATGTCTGGGAGCCCGCAGTTTTCATCCTTACATACAAACCGACTTTTGTGAATTCCAAATGGAACTGATTACACCAGTAGCCAAAAGTACGACGGAGGCCCGACGCTTTCTTGGCGCGATAACTGATGTAGCAGGACGAAGCATTGCAACTGACGAGGTGCTGTGGCCATTGAGTATGCCACCACGACTTAAAGCCGAGGAAATTCAAGTAGCGCAACTCGAGAACGACTTCGAAAGACATTATCGGAACTACTTGGCAGAGAAGTACGGCACCAAATTGCAGGCGATTAGTGGAATTCATTACAATATGGAACTTGGGAAGGACTTGGTTGAGGCGCTTTTTCAAGAGTCAGATCAGACTGACATGATCGCATTTAAAAACGCTCTGTATCTCAAGCTCGCCCAGAACTATTTGAGGTATCGGTGGGTCATTACTTACCTGTTTGGAGCAAGTCCCATTGCAGAACAAGGATTCTTTGACCAAGAAGTGCCGGAGCCTATGCGCTCTTTCCGCAACTCCGACCACGGCTACGTTAACAAGGAAGAGATACAGGTAAGCTTTGTATCCCTTGAAGACTATGTCTCCGCGATCGAGACCTACATCGAGCAGGGTGACCTTATAGCCGAGAAAGAGTTTTACTCAGCCGTGCGCTTTAGAGGACAAAAAGTCAATCGCTCCTTCCTTGATAAGGGTATAACTTATCTGGAGTTCAGAAACTTTGACTTGAACCCATTTGAGAGAATAGGCATCAGTCAAACCACTATGGATACCGTTCACTTGCTCATACTGGCCTTTCTCTGGTTGGATAGTCCGGAAAACGTGGACCAGGCCCTTGCCCAGGGACACGCGCTTAACGAGAAAATAGCCCTCTCCCATCCATTGGAGCCCCTCCCTTCAGAGGCCAAGACACAGGACATCGTGACCGCACTCGACCAGCTCGTACAGCACTTCGGATTGGGAGATTACCACCAGGATCTCGTGAAACAAGTGAAAGCGGCGTTTGCTGATCCGAATCAAACCCTGTCAGCTCAACTTCTTCCTTATATTAAGGACAAGTCACTCGCAGAATTCGCTCTCAATAAGGCACTCGCATATCATGACTATGACTGGACCGCTCACTACGCCCTTAAAGGTTACGAAGAAATGGAGCTCAGTACGCAGATGCTGCTCTTTGATGCTATCCAGAAAGGAATACACTTCGAGATACTCGATGAGCAAGATCAGTTCTTGAAGCTGTGGCATCAAGATCACGTAGAATATGTTAAAAACGGTAATATGACCAGCAAGGATAACTATGTAGTACCTCTCGCAATGGCCAACAAGACTGTTACTAAGAAAATTCTTGCTGACGCTGGGTTCCCTGTTCCGTCCGGGGACGAATTTACTAGCTTGGAGGAGGGACTGGCCTACTACCCGCTTATTAAAGATAAGCAAATTGTAGTAAAGCCAAAGAGCACGAATTTCGGCTTGGGTATCAGCATCTTCCAAGAACCCGCCAGTCTCGACAATTATCAAAAAGCATTGGAAATAGCATTTGCGGAGGACACTAGTGTGCTCGTCGAAGAATTCATTCCAGGCACGGAATACCGATTCTTCATTTTGGACGGACGCTGTGAGGCAGTCCTTTTGAGGGTAGCTGCCAATGTAATAGGGGACGGGAAACACACAATCAGAGAGTTGGTAGCGCAGAAAAACGCAAATCCCCTGCGCGGTAGGGATCATAGATCACCCTTGGAAATCATAGAGCTTGGGGACATAGAGCAACTCATGCTGGCACAGCAGGGTTACACTCCAGATGACATCCTGCCAGAGGGTAAGAAAGTGAATTTGAGGCGGAACAGCAATATTAGTACTGGGGGAGACTCCATAGACGTCACAGAAACAATGGATAGCT CTTATCAAGAACTTGCAGCAGCGATGGCTACCAGTATGGGGGCATGGGCCTGTGGAGTTGATCTGATTATACCCGACGAAACGCAGATTGCCACAAAGGAAAATCCACATTGCACGTGTATTGAACTTAACTTCAACCCCTCCATGTACATGCATACATACTGCGCTGAGGGGCCGGGGCAGGCAATTACAACCAAAATACTCGACAAACTCTTCCCGGAGATCGTGGCCGGACAAACTTAAGCTACGTAAATTCCGCCC

Human codon-optimized Cyto-GshF was custom synthesized by IDT and directly assembled into pMXS-puro linearized with BamHI and NotI with a Gibson reaction.

To add the MTS, the GshF gene was divided into three fragments: MTS, middle, and C-terminal, with 5’ and 3’ regions designed to overlap with the preceding or proceeding fragment or the pMXS vector. Each fragment was custom synthesized by Twist Bioscience. Fragments were PCR amplifed and gel extracted with a Zymo gel extraction kit before being assembled into pMXS-puro linearized with BamHI and NotI with a 4-component Gibson reaction.

Primers:

5’-GCCGGATCTAGCTAGTTAATTAAGccacc-3’; MTS_Fwd
5’-CCATTTGGAATTCACAAAAGTCGGTTTGT-3’; MTS_Rev:
5’-ACAAACCGACTTTTGTGAATTCCAAATGG-3’; Mid_GSHf_Fwd
5’-GGAAGAAGTTGAGCTGACAGGGTTTGATT-3’; Mid_GSHf_rev
5’-ATGAAACCCTGTCAGCTGAACTTC-3’; C-GSHf_Fwd
5’-GGGCGGAATTTACGTAGCTTAAGT-3’;C-GSHf_Rev

#### List of other constructs in the manuscript

PMXS_IRES_BLAST_Hu_SLC25A39; PMXS_IRES_BLAST_Hu_SLC25A39_D226A mutant; PMXS_IRES_BLAST_Hu_SLC25A39_K329A mutant; PMXS_IRES_BLAST_Hu_SLC25A40; PMXS_IRES_PURO_mito-Gshf; PMXS_IRES_PURO_cyto-Gshf; PMXS_IRES_GFP_cyto-Gshf; PMXS_IRES_GFP_mito-Gshf; PMXS_IRES_BLAST_mito-Gshf; PMXS_IRES_BLAST_cyto-Gshf; lentiCRISPR-v1_sg4_Hu_SLC25A39 (GFP); lentiCRISPR-v2_sg1_Hu_SLC25A40 (PURO); lentiCRISPR-v2_sg4_Hu_GCLC (PURO).

### Immunoblotting

Cell pellets were washed twice with ice-cold PBS prior to lysis in RIPA buffer (10 mM Tris-Cl pH 7.5, 150 mM NaCl, 1 mM EDTA, 1% Triton X-100, 0.1% SDS) supplemented with protease inhibitors Sigma-Aldrich, 11836170001) and phosphatase inhibitors (Roche, 04906837001). Each lysate was sonicated and, after centrifugation for 5 min at 4°C and 20,000 x g, supernatants were collected. Sample protein concentrations were determined by using Pierce BCA Protein Assay Kit (Thermo Scientific) with bovine serum albumin as a protein standard. Samples were resolved on 12% or 10-20% SDS-PAGE gels (Invitrogen) and analyzed by standard immunoblotting protocol.

### Cell proliferation assays

For cell proliferation assays with cell counting, cells were plated in triplicates in 24-well plates at 10000 cells/well. Cells were collected after 4 days and counted by Z2 Coulter Counter (Beckman). For cell proliferation assays with fold change in cell number, cells were cultured in replicates of 3 in 96-well plates at 1000 (suspension) or 150 (adherent) cells per well in 200 uL RPMI or DMEM base media under the conditions described in each experiment, and a separate group of 3 wells was also plated for each cell line with no treatment for an initial time point. Adherent cells were allowed 1 day to adhere. Initially (untreated cells for initial time point) or after 4 days (with varying treatment conditions), 40 μL of Cell Titer Glo reagent (Promega) was added to each well, mixed briefly, and the luminescence read on a luminometer (Molecular Devices). For each well, the fold change in luminescence relative to the initial luminescence was measured and reported in a log_2_ scale as the number of cell doublings. Cell culture images were taken using a Primovert microscope (Zeiss).

### Whole cell and mitochondrial proteomics

After the indicated culture conditions, cell pellets were processed as per immunoblot protocol and lysed with lysis buffer (50 mM Tris-Cl pH 7.5, 150 mM NaCl, 1 mM EDTA, 2% Triton X-100) supplemented with protease inhibitors and phosphatase inhibitors (Roche).

#### Digestion

Samples were dried and dissolved in 8M urea, 50mM triethylammonium bicarbonate (TEAB), 10 mM dithiothreitol (DTT) and disulfide bonds were reduced for 1 hour at room temperature. Alkylation was performed using iodoacetamide (IAA) for 1 hour at room temperature in the dark. Proteins were precipitated by Wessel/Flügge extraction^37^ and pellets were dissolved in 100mM TEAB with endopeptidase LysC (2% w/w, enzyme/substrate) and incubated at 37°C for 2-3 hours. Sequencing grade modified trypsin (2 % w/w, enzyme/substrate) was added, and digestion proceeded overnight.

#### Labeling and fractionation

Peptide solutions were labeled with 270 μg aliquots of TMTpro (Thermo Scientific) for 1 hour at room temperature and subsequently quenched with hydroxylamine for 15 minutes. An aliquot from each sample was combined for a ratio check, according to which the samples were mixed. The pooled sample was purified using a high-capacity reverse phase cartridge (Oasis HLB, Waters) and the eluate was fractionated using high pH reverse phase spin columns (Pierce) according to manufacturer specifications, yielding 8 fractions.

#### LC-MS/MS

Fractionated peptides were analyzed using an Easy-nLC 1200 HPLC equipped with a 250mm*75μm Easyspray column connected to a Fusion Lumos mass spectrometer (all Thermo Scientific) operating in synchronous precursor selection (SPS)-MS3 mode^38^ (10 SPS events). Solvent A was 0.1% formic acid in water and solvent B was 80% acetonitrile, 0.1% formic acid in water. Peptides from the mitochondrial-IP were separated across a 90-minute linear gradient and peptides from the whole-cell lysate were separated across a 120-minute linear gradient going from 7 to 33% B solvent at 300nL/minute. Precursors were fragmented by CID (35% CE) and MS2 ions were measured in the ion trap. MS2 ions were fragmented by HCD (65% CE) and MS3 reporter ions were measured in the orbitrap at 50K resolution.

#### Data analysis

Raw files were searched through Proteome Discoverer v.2.3 (Thermo Scientific) and spectra were queried against the human proteome (database downloaded from uniprot.org on 02/12/2019, containing 73662 sequences) using Sequest HT with a 1 % false discovery rate (FDR) applied. Oxidation of M was applied as a variable modification and carbamylation of C was applied as a static modification. A maximum isolation interference of 50% was allowed and 80% matching SPS ions were required. Protein abundance values were used for further statistical analysis. A complete list of identified peptides is provided in **Extended Data Table 3**.

#### Statistical analysis

Subsequent statistical analysis was performed within the Perseus framework^39^. All values were log2 transformed and normalized to the median intensity within each sample. An FDR-corrected t-test (q=0.05) was used to test for significance between sample groups.

For mitochondrial proteomics, mitochondria were rapidly immunopurified from HeLa cells expressing a mitotag (3xHA-OMP25-mCherry) grown in indicated cell culture conditions, according to the rapid mitochondrial purification for metabolite profiling protocol. At the final step after the third KPBS wash, the entire bead volume was lysed in 70□μl of lysis buffer (50 mM Tris-HCl, pH 7.4, 150 mM NaCl, 1 mM EDTA, 1% Triton X-100 with protease inhibitors (Roche). Approximately 50 □μg of protein extracted from mitochondria were submitted for proteomics. Peptides were analyzed similarly to whole cell proteomics and a complete list of identified peptides is provided in **Extended Data Table 1**. Due to high isolation interference, the SLC25A39 peptide TMTpro-GLFAGFLPR (m/z 641.3848) underwent a targeted MS3 strategy. MS2 fragmentation was performed as described in the method section, but only the singly-charged b7-ion (m/z 1010.5) was targeted for MS3 fragmentation. Due to the low reporter ion intensities resulting from fragmentation of a low-abundant peptide and selecting only one fragment, the reporter ion intensities were summed across the chromatographic elution profile of the peptide. Missing values were imputed with random low-abundant numbers from a normal distribution and the result was incorporated into the volcano plot in **Figure 1C**.

### Polar metabolite profiling

After the indicated culture conditions, cells were washed with 1 mL of cold 0.9% NaCl. Polar metabolites were extracted in 0.5 mL of cold 80% methanol containing internal standards (Cambridge Isotope Laboratories). After extraction, samples were nitrogen-dried and stored at −80C until analysis by LC-MS. Analysis was conducted on a QExactive benchtop orbitrap mass spectrometer equipped with an Ion Max source and a HESI II probe, which was coupled to a Dionex UltiMate 3000 UPLC system (Thermo Fisher Scientific). External mass calibration was performed using the standard calibration mixture every 7 days.

Dried polar samples were resuspended in 100 μL water and 2 μL were injected into a ZIC-pHILIC 150 x 2.1 mm (5 μm particle size) column (EMD Millipore). Chromatographic separation was achieved using the following conditions: Buffer A was 20 mM ammonium carbonate, 0.1% ammonium hydroxide; buffer B was acetonitrile. The column oven and autosampler tray were held at 25°C and 4°C, respectively. The chromatographic gradient was run at a flow rate of 0.150 mL/min as follows: 0–20 min.: linear gradient from 80% to 20% B; 20–20.5 min.: linear gradient from 20% to 80% B; 20.5–28 min.: hold at 80% B. The mass spectrometer was operated in full-scan, polarity switching mode with the spray voltage set to 3.0 kV, the heated capillary held at 275°C, and the HESI probe held at 350°C. The sheath gas flow was set to 40 units, the auxiliary gas flow was set to 15 units, and the sweep gas flow was set to 1 unit. The MS data acquisition was performed in a range of 70–1000 m/z, with the resolution set at 70,000, the AGC target at 10e6, and the maximum injection time at 20 msec. Relative quantitation of polar metabolites was performed with XCalibur QuanBrowser 2.2 (Thermo Fisher Scientific) using a 5 ppm mass tolerance and referencing an in-house library of chemical standards. Metabolite levels were normalized to the total protein amount for each condition.

### Rapid mitochondrial purification for metabolite profiling

Mitochondria were purified from HEK-293T, HeLa, or K562 cells expressing 3xHA-OMP25-mCherry (mitochondrial isolation) or 3xMyc-OMP25-mCherry (background control) according to a previously described protocol^40^. Briefly, 25-30 million cells were collected and washed twice with cold saline (0.9% NaCl), scraped into 1 □ml of cold KPBS, and pelleted via centrifugation at 1,000g for 1.5□min at 4□°C. Cells were resuspended in 1 □ml of KPBS, 10□μl of cells were transferred into 40□ml of 1% Triton lysis buffer for a whole-cell protein sample and 10□μl of cells were transferred into 50□ml of 80% methanol, containing heavy labeled amino acid standards, for direct extraction of whole-cell metabolites. With one set of 20 strokes and another set of 10 strokes, the remaining sample was homogenized using a 2-ml homogenizer. After centrifugation, the homogenate was incubated with 200□ml of KPBS pre-washed anti-HA magnetic beads (Thermo Scientific Pierce 88837) on a rotator shaker for 5□min at 4□°C. Beads were washed three times in cold KPBS, then 10% of bead volume was lysed with 1% Triton buffer for protein extracts and the remaining 90% was extracted in 80% methanol containing heavy labeled amino acid standards (Cambridge Isotope Labs) on a rotator shaker for 10□min at 4□°C.

Samples were spun down at 20,000g to remove potential cellular debris or bead contamination. Samples were subjected to LC-MS polar metabolite profiling without drying. Data were normalized by CS protein levels (western blot) or NAD+ abundance (see details in figure legends).

### Glutathione uptake assay in immunopurified mitochondria

Mitochondria were purified from HEK-293T cells expressing 3xHA-OMP25-mCherry (mitochondrial isolation) or 3xMyc-OMP25-mCherry (background control) according to a previously described protocol^40^. Briefly, 30 million cells were collected and washed twice with cold saline (0.9% NaCl), scraped into 1□ml of cold KPBS, and pelleted via centrifugation at 1,000g for 1.5□min at 4□°C. Cells were resuspended in 1□ml of KPBS and 10□μl of cells were transferred into 40□ml of 1% Triton lysis buffer for a whole-cell protein sample. With one set of 20 strokes and another set of 10 strokes, the remaining sample was homogenized using a 2-ml homogenizer. After centrifugation, the homogenate was incubated with 200□ml of KPBS pre-washed anti-HA magnetic beads on a rotator shaker for 5□min at 4□°C. Following incubation, beads were washed once with cold KPBS (pH=7.25) before being incubated in 200 μl transport buffer (KPBS+10 mM HEPES + 0.5 mM EGTA) at room temperature containing indicated doses of [^13^C_2_,^15^N]-GSH (Cambridge Isotope Laboratory, CNLM-6245-50) for the indicated times. It is important to maintain the pH between 7.3 and 7.4 by adding 0-2 μl 2M KOH per ml of transport buffer. Uptake was stopped with the addition of cold KPBS and beads were subsequently washed three more times in cold KPBS. Following the third wash, 10% of the bead volume was lysed with 1% Triton buffer for protein extracts and the remaining 90% was extracted in 80% methanol containing heavy labeled amino acid standards on a rotator shaker for 10 min at 4 °C. Samples were spun down at 20,000g and the uptake of [^13^C_2_,^15^N]-GSH into mitochondria is measured by LC-MS.

### Aconitase assay

Assay was carried out with the Aconitase Activity Assay Kit (Abcam ab109712). Cell lines were generated as indicated and each condition was collected in triplicates of 1,000,000 cells each. Cell pellets were washed with ice-cold PBS prior to lysis in 1 mL aconitase preservation solution with detergent. Lysates were kept on ice for 30 mins and centrifuged for 10 min at 4°C and 20,000 x g. 50 μL of the supernatant was transferred to a well on the assay plate and 200 μL of assay buffer was added to each well. Plate was read on a plate reader (Molecular Devices) according to manufacturer’s protocol with a kinetic program of 45 second intervals.

### CRISPR-based genetic screens

The metabolism-focused human sgRNA library was designed and screens performed as previously described^41^. Oligonucleotides for sgRNAs were synthesized by CustomArray Inc. and amplified by PCR. A complete list of differential gene scores for each screen is provided in **Extended Data Table 2**. Briefly, plasmid pool was used to generate a lentiviral library, which was transfected into HEK-293T cells and used to generate viral supernatant as described above. SLC25A39 knockout cells expressing a vector control or SLC25A39 cDNA, in both Jurkat and HEK-293T, were infected at a multiplicity of infection of 0.7 and selected with puromycin. An initial sample of 30 million cells was harvested and infected cells were cultured for ~14 population doublings. Final samples of 30 million cells were collected. DNA was extracted (DNeasy Blood and Tissue kit, Qiagen). sgRNA inserts were PCR-amplified and barcoded by PCR, using primers unique to each condition. PCR amplicons were purified and sequenced (NextSeq 500, Illumina). Screens were analyzed using Python (v.2.7.13), R (v.3.3.2) and Unix (v.4.10.0-37-generic x86_64). The gene score for each gene was defined as the median log2 fold change in the abundance of each sgRNA targeting that gene. All gene scores for each screen are provided in **Extended Data Table 2**.

### Immunofluorescence

For SLC25A39 localization immunofluorescence assays, 100,000 Hela cells were seeded onto coverslips. 24 hours after plating, cells were treated with vehicle or BSO (1mM). After 24 hours of treatment, cells were fixed for 10 minutes at −20 °C with ice-cold acetone. After three PBS washes, cells were blocked with 0.5% BSA in PBS, and incubated with the anti-SLC25A39 (Proteintech, 14693-1-AP, 1:200) and anti-ATP5A (Thermo Fisher Scientific, 439800, 1:200) primary antibodies in blocking solution for one hour. Cells were washed three times with PBS and incubated with a donkey anti-mouse Alexa Fluor 568 (ThermoFisher Scientific, A10037) and donkey anti-rabbit Alexa Fluor 488 (ThermoFisher Scientific, A21206) secondary antibodies (1:400) for one hour in the dark. Cells were washed three times with PBS, and incubated with a 200 nM solution of DAPI in the dark. Cells were washed three final times, and the coverslip was mounted onto slides with Prolong Gold antifade mounting media (Invitrogen). Confocal images were acquired with a Zeiss inverted LSM 789 laser scanning confocal microscope (Zeiss) using a 100x/1.4 DIC Plan-Apochromat oil immersion objective. Four representative fields were captured for each condition under identical exposure times. Images were obtained with excitation and emission wavelengths as follows: DAPI 420-498, Alexa Fluor 568 578-650, Alexa Fluor 488 498-552. The images are 512 x 512 pixels with a pixel depth of 12-bit, a pixel size of 0.08 μM per pixel, a dwell time of 6.30μs, a pinhole size of 60 (1Airy unit), and a line averaging of 1.

For mito- and cyto-gshf localization immunofluorescence assays, 100,000 293T cells were seeded onto coverslips coated with poly-D-lysine. After 24 hours, the cells were fixed at room temperature for 10 minutes with 4% paraformaldehyde in PBS. After three PBS washes, cells were permeabilized with 0.1% Triton-X for 10 minutes, shaking. After three additional PBS washes, cells were blocked in 5% normal donkey serum (NDS) for 1h at room temperature, shaking. The coverslips were subsequently incubated with anti-Flag-M2 (Sigma, F3165, 1:400) and anti-citrate synthase (Cell Signaling, 14309S, 1:200) for 16h at 4 °C before washing 3 times with PBS. Coverslips were then incubated with secondary (Alexa Flour 568 anti-mouse and Alexa Flour 488 anti-rabbit) 1:500 in 5% NDS for 1 h at room temperature in the dark and washed 3 times with PBS. Cells were incubated with a solution of 200nM in the dark before and washed an additional 3 times with PBS. Finally, coverslips were mounted onto slides with Prolong Gold Antifade mounting media (Invitrogen). Images were obtained with a Nikon A1R MP+ multiphoton confocal microscope (Nikon) using a 60X/1.4 DIC Plan-Apochromat oil immersion objective. Images were obtained with excitation wavelengths as follows: DAPI 425-475, Alexa Fluor 488 500-550, Alexa Fluor 568 570-620. The images are 1024 x 1024 pixels with a pixel depth of 12-bit, a pixel size of 0.07μm, a dwell time of 0.5s, a pinhole size of 39.59μM (1.2 Airy units), and a line averaging of 4.

### Generation of Slc25a39 KO and conditional KO mice

All animal studies and procedures were conducted according to a protocol approved by the Institutional Animal Care and Use Committee (IACUC) at the Rockefeller University. All mice were maintained on a standard light-dark cycle with food and water ad libitum. All treatment studies were randomized and injections were performed by blinded investigators.

CRISPR guide RNAs were designed using CHIRPOR.org^42^ and were used as two-part synthetic crRNA and tracrRNA (Alt-RTM CRISPR guide RNA, Integrated DNA Technologies, Inc). Cas9 protein, crRNA, and tracrRNA were assembled to ctRNP using protocols described previously^43^. All guide RNAs were validated in mouse embryos for their efficiency of generating indels on target genomic sequence, and two gRNAs targeting intron 1 and intron 2 were selected for preparing the microinjection mix. sg1. AATGTGCTCACTTTGCCGAGGGG; sg2. TCTATGATCATGGGTAACCCAGG

The Slc25a39-flox-E2-pUC57 plasmid was constructed to include a floxed Slc25a39 exon2 and 400 bp of homologous sequence on each side. To prepare the long single-stranded editing donor DNA (lssDNA), the 1.5 kb insert was amplified from slc25a39-flox-E2-Puc57 and processed to lssDNA using exonucleolytic hydrolysis of its 5’-phosphorylated DNA strand as described previously^44^.

The final injection mix was made of 0.6μM of each guide RNA (crRNA + tracrRNA), 0.3μM of Cas9 protein, and 10ng/ μl of ssDNA according to protocols described. The injection mix was then delivered to 0.5 days of fertilized C57Black/6 mouse embryos using well-established pronuclear injection and surgical protocols^43^. All animal experiments followed the approved IACUC protocols.

### Histology and Immunohistochemistry

Paraformaldehyde-fixed and paraffin-embedded tissues were deparaffinized and rehydrated, then incubated with anti-Ter119 antibody (Biolegend, 116201, 1:200) at 4°C overnight and with secondary Goat anti-Rat IgG (VECTOR, PI-9401) for 1 hour at room temperature. The reaction was developed with diaminobenzidine (DAB) staining (VECTOR, SK-4100). Slides were then counterstained in nuclear fast red. Stained slides were analyzed in a blinded fashion.

### Cell Profiling of Fetal Liver Cells

E12.5 fetal liver cells were resuspended in 2% FBS and 10 mM glucose in PBS. For analysis of erythroid progenitor cells, fetal liver cells were stained with PE-anti mouseTer119 (Biolegend, 116207, 1:200), PE/Cy7-anti CD44 (Biolegend, 103029, 1:200) and DAPI. For analysis of stem cells, fetal liver cells were stained with the following antibodies at 1:100 dilution: PECy5-B220 (ebioscience 15-0452-83), PECy5-CD3 (ebioscience 15-0031-83), PECy5-Mac1 (ebioscience 15-00112-83), PECy5-Gr1 (ebioscience 15-5931-82), PECy5-Ter119 (ebioscience 15-5921-83), PECy5-CD4 (ebioscience 15-0041-83), PECy5-CD8 (ebioscience 15-0081-83), Pacific Blue-Sca1 (biolegend 122520), APCCy7-Ckit (biolegend 105826), FITC-CD34 (ebioscience 11-0341-85), PECy7-cd16/cd32 (BD pharmingen 560829); myeloid cells, PECy5-Ter119 (ebioscience 15-5921-83), FITC-CD71 (ebioscience 11-0711-82), PE-CD41 (BD pharmingen 558040), Pacific Blue-Mac1 (biolegend 101224), APCCy7-GR1 (biolegend 108424); lymphoid cells, Pacific Blue-CD3 (ebioscience 48-0031-82), APC-IgM (biolegend 406509), PECy7-B220 (BD pharmingen 552772). Flow cytometry was performed using the LSRII (Beckton Dickinson) and Attune NxT Flow Cytometer. Data were analyzed using FlowJo software (Beckton Dickinson) and FCS Express 7.

### Gene Ontology, co-evolution analysis and homology modeling

Genes with phylogenetic profiles similar to SLC25A40 was determined using Phylogene^45^. Phylogenetic tree for SLC25A39 is constructed using Treefam. Gene ontology analysis for proteomics results has been performed with PANTHER and GORILLA^46^.

A homology model of human SLC25A39 was generated using the SWISS-MODEL server homology modelling pipeline with the bovine ATP-ADP carrier (PDB: 1OKC) as a scaffold ^47,48^. Substrate-binding residues in human SLC25A39 were predicted based on biochemical and genetic analyses of the related ATP-ADP carrier using the ClustalW algorithm in MegAlign Pro 16 (DNASTAR)^19,49^. Structural depictions of the SLC25A39 homology model were prepared in Chimera 1.14^50^.

### RNA Extraction and Real-time PCR

RNA was extracted from 293T using the Qiagen RNeasy mini kit according to the manufacturer’s instructions. 1 ug of RNA was reverse transcribed using the Superscript III Reverse Transcriptase kit (Invitrogen) according to the manufacturer’s instructions. Quantitative real-time PCR (qPCR) was performed using SYBR green master mix and *β-ACTIN* was used as a control. The primer sequences are as below:

*β-ACTIN F*: CATGTACGTTGCTATCCAGGC; *R*: CTCCTTAATGTCACGCACGAT
*SLC25A39 F: TGCCCTTCTCAGCCCTGTA; R: GGTTCACTCTCACAGCCTCC*
*GCLM F*: TGTCTTGGAATGCACTGTATCTC; *R*: CCCAGTAAGGCTGTAAATGCTC

### Cell Mixing Competition Assays

A small competition sgRNA library targeting SLC25A40 and TXNRD2 (5 sgRNAs each) in addition to 5 sgRNAs controls was designed from a larger focused sgRNA metabolism library. Oligos (IDT) were cloned into lentiCRISPR-v1 (controls) or lentiCRISPR-v2 (SLC25A40 and TXNRD2) using T4 ligase (NEB). Ligation products were cloned into Escherichia coli (NEB) and plasmids were isolated by Miniprep (Qiagen). This plasmid pool was used to generate a lentiviral library, which was transfected into HEK-293T cells and used to generate viral supernatant as described above. HEK-293T SLC25A39 knockout cells expressing a vector control or SLC25A39 cDNA were infected at a multiplicity of infection of 0.7 and selected with puromycin. An initial sample of 30 million cells was harvested and infected cells were cultured for ~14 population doublings in either standard cell culture conditions or treated with 20□μM BSO. Final samples of 30 million cells were collected. DNA was extracted (DNeasy Blood and Tissue kit, Qiagen). sgRNA inserts were PCR-amplified and barcoded by PCR, using primers unique to each condition. PCR amplicons were purified and sequenced (NextSeq 500, Illumina). Screens were analyzed using Python (v.2.7.13), R (v.3.3.2) and Unix (v.4.10.0-37-generic x86_64). The gene score for each gene was defined as the median log2 fold change in the abundance of each sgRNA targeting that gene.

Oligo sequences for competition library: Human SLC25A40 sg1 F: caccGGTTGGATTTCCCTTTGGAG, sg1 R: aaacCTCCAAAGGGAAATCCAACC, sg5 F: caccgAAGAGGGAGGCAACAAACTA, sg5 R: aaacTAGTTTGTTGCCTCCCTCTTc, sg6 F: caccgAAGAAGCCAGGAAATTTCCA, sg6 R: aaacTGGAAATTTCCTGGCTTCTTc, sg7 F: caccGGTACATCTCTAAGAACAGT, sg7 R: aaacACTGTTCTTAGAGATGTACC, sg8 F: caccgCTATGTGTCTGTGAAGAGGG, sg8 R: aaacCCCTCTTCACAGACACATAGc, Human TXNRD2 sg2 F: caccgTAAACCACTGGAGTTCACGG, sg2 R: aaacCCGTGAACTCCAGTGGTTTAc, sg3 F: caccGATTAGGAGGGCGCTTCCGG, sg3 R: aaacCCGGAAGCGCCCTCCTAATC, sg6 F: caccgCAAGAAGCTGATGCACCAGG, sg6 R: aaacCCTGGTGCATCAGCTTCTTGc, sg7 F: caccGCACTACCTGGTCGAAGCCG, sg7 R: aaacCGGCTTCGACCAGGTAGTGC, sg8 F: caccGGAGGACAGCACCACCGGCA, sg8 R: aaacTGCCGGTGGTGCTGTCCTCC, Control sg5 F: GCGGATTAGAGGTAATGCGG, sg5 R: CCGCATTACCTCTAATCCGC, sg6 F: GGAGCCATGGTAGAGCGTAT, sg6 R: ATACGCTCTACCATGGCTCC, sg11 F: GGCGCTCGATTAAGACTCAG, sg11 R: CTGAGTCTTAATCGAGCGCC, sg12 F: GTTACGCGTGCTGTCGCTTC, sg12 R: GAAGCGACAGCACGCGTAAC, sg13 F: GTTCCTTCTGCAGGCACGCT, sg13 R: AGCGTGCCTGCAGAAGGAAC

#### Analysis of transcriptome-wide association studies (TWAS) data

We analyzed transcriptome-wide association studies (TWAS) results on 4,000 traits for SLC25A39 in the UK Biobank^12^ (n=361,194) using the summary-statistics-based, joint-tissue PrediXcan^51,52^. Application of PrediXcan to a large-scale biobank has been shown to enable discovery of gene-level effects on the medical phenome^53^. Here, for the gene expression models, we leveraged the GTEx V8^54^ transcriptome panel. This approach regresses each phenotype on the principal components of the estimated genetically-determined SLC25A39 expression data across the 49 tissues (Suppl. Table X). We applied Bonferroni adjustment for multiple testing correction (p_adjusted<0.05). We report the most significant results (i.e., for hematological traits) from the unbiased analysis across the entire phenomic dataset.

#### Statistics and reproducibility

GraphPad PRISM 7 and Microsoft Excel 15.21.1 software were used for statistical analysis. XCalibur QuanBrowser 2.2 (Thermo Fisher Scientific) was used for metabolomic analyses and ImageJ (NIH) for image analysis. Error bars, *P* values and statistical tests are reported in the figure captions. All experiments (except full metabolite profiling experiments and proteomics, which were done once) were performed at least two times with similar results. Both technical and biological replicates were reliably reproduced. Comparison of two mean values was evaluated by two-tailed unpaired *t*-test. Comparison of multiple mean values was evaluated by One-way ANOVA followed by post-hoc Bonferroni test. Comparison of multiple mean values under different conditions was evaluated by Two-way ANOVA. Statistical significance was determined by two-tailed unpaired *t*-test.

#### Data availability

All data supporting the findings of this study are available from the corresponding authors on reasonable request.

